# Century-long timelines of herbarium genomes predict plant stomatal response to climate change

**DOI:** 10.1101/2022.10.23.513440

**Authors:** Patricia L.M. Lang, Joel M. Erberich, Lua Lopez, Clemens L. Weiß, Gabriel Amador, Hannah F. Fung, Sergio M. Latorre, Jesse R. Lasky, Hernán A. Burbano, Moisés Expósito-Alonso, Dominique Bergmann

**Author notes:** **Author contributions:** PLML conceived and designed the project with input from MEA, HAB and DB; PLML designed and performed analyses with contributions from MEA, CLW and JE; LL, JRL, SL and HAB contributed data; SML helped with historical sample collection and processing; HFF contributed microscopy; GA and HAB contributed to discussion of results; MEA provided computational resources; DCB supervised research; PLML and JE wrote the manuscript with input from all authors.

## Abstract

Dissecting plant responses to the environment is key to understanding if and how plants adapt to anthropogenic climate change. Stomata, plants’ pores for gas exchange, are expected to decrease in density following increased CO_2_ concentrations, a trend already observed in multiple plant species. However, it is unclear if such responses are based on genetic changes and evolutionary adaptation. Here we make use of extensive knowledge of 43 genes in the stomatal development pathway and newly generated genome information of 191 *A. thaliana* historical herbarium specimens collected over the last 193 years to directly link genetic variation with climate change. While we find that the essential transcription factors SPCH, MUTE and FAMA, central to stomatal development, are under strong evolutionary constraints, several regulators of stomatal development show signs of local adaptation in contemporary samples from different geographic regions. We then develop a polygenic score based on known effects of gene knock-out on stomatal development that recovers a classic pattern of stomatal density decrease over the last centuries without requiring direct phenotype observation of historical samples. This approach combining historical genomics with functional experimental knowledge could allow further investigations of how different, even in historical samples unmeasurable, cellular plant phenotypes have already responded to climate change through adaptive evolution.

**One sentence summary:** Using a molecular-knowledge based genetic phenotype proxy, historical whole-genome *A. thaliana* timelines compared with contemporary data indicate a shift of stomatal density following climate-associated predictions.

## Introduction

Ongoing drastic increases in CO_2_ concentrations, and the resulting changes in global temperatures and droughts are altering our environment (Pörtner et al., n.d.). Dissecting if and how plants respond to this will be key to understanding plants’ potential for adaptation to climate change, and to developing strategies to increase their chances of survival. Several phenotypic trends observed in large numbers of plant species and across continents are being reported, including the acceleration of flowering (Panchen et al. 2012; Primack et al. 2004) and other life history events (Parmesan and Yohe 2003), the increase in photosynthesis (Walker et al. 2021), and the decrease in the number of stomatal pores in plant leaves (Woodward 1987). However, it has thus far been difficult to resolve whether these trends reflect plastic phenotypic changes or result from evolutionary genetic change (Parmesan 2006). Stomata are one of the plant structures that are most directly relevant to multi-factorial climatic changes, as these surface pores, essential for survival and productivity, regulate plants’ water-use efficiency (WUE): the ratio between CO_2_ uptake for photosynthesis, and the release of O_2_ and loss of water vapor with transpiration (reviewed in (Han, Kwak, and Qi 2021)). Optimized WUE to environmental conditions ensures maximal plant growth and efficient cooling through transpiration while minimizing water loss. It is partially fine-tuned through variation in stomatal size and density (i.e. the amount of stomata per plant surface area; (Bertolino, Caine, and Gray 2019; Faralli, Matthews, and Lawson 2019; Berry, Beerling, and Franks 2010)). This variation can represent temporary, plastic responses, or result from evolutionary genetic change over generations that generates local adaptation (Vinton et al. 2022; Bay et al. 2017; Gienapp et al. 2008).

A powerful way to differentiate between plastic and evolutionary plant responses to climate is to study different populations of a single species collected across geographic climatic gradients in combination with genetic analyses (Clausen, Keck, and Hiesey 1941). Both stomatal size and density correlate with climate gradients, indicating a potential genetic basis for their climate responses (Dittberner et al. 2018). In *Arabidopsis*, lower stomatal density typically follows increasing CO_2_ and temperature, while higher stomatal densities and drought-adjusted higher WUE result from decreased humidity. Stomatal size behaves anti-correlated to stomatal density (Crawford et al. 2012; Vile et al. 2012; Yan, Zhong, and Shangguan 2017; Lau et al. 2018; Dittberner et al. 2018). The last ~200 years of anthropogenic global change have further altered this spatiotemporal plant variation in complex unknown ways. Collections of pressed and dried specimens are witnesses of plant responses to the CO_2_-increase from ~280 parts per million in 1750 to 419 ppm in 2022 (August measurement, see trends 2 at www.climate.gov and climate.nasa.gov), or the current global maximum temperature anomaly of ca. +1°C (climate.nasa.gov), and with that track adaptation to climate change as it happens (reviewed in (Lang et al. 2018)).

Plant responses themselves can help inferring historical climate trends. Measurements of fossilized stomatal densities indicate atmospheric CO_2_ changes over geological time (Beerling and Chaloner 1992; Van Der Burgh et al. 1993; Beerling and Chaloner 1993; McElwain and Chaloner 1995), and over the recent anthropogenic climate change, decreases in stomatal densities preserved in herbaria already reflect the industrialization-related increases in atmospheric CO_2_ (Woodward, 1987). Such analyses have thus far never extended beyond phenotypic quantification to also assess genetic changes underlying these potentially adaptive responses, mainly because fossil records lack quantifiable DNA. Now, sequencing of herbarium specimens can address this gap by directly exploring joint timelines of phenotypic and genotypic responses to climate change (e.g. (Yoshida et al. 2013; Lang et al. 2020; Latorre et al. 2020; Kistler et al. 2020)).

Stomata are a unique system to use such timelines to understand plants’ adaptive potential and its mechanisms, as their genetic pathway is dissected in minute detail in *A. thaliana* (e.g. reviewed in (L. R. Lee and Bergmann 2019; Kinoshita, Toh, and Torii 2021; Han, Kwak, and Qi 2021)): From the sequential interactions of indispensable transcription factors that regulate cell production, fate and patterning, to external regulators that fine-tune stomatal development in response to environmental and physiological stimuli. This knowledge provides a crucial advantage when investigating the genetic basis of potential stomata involvement in climate adaptation. It allows selective study of genetic variants in already-validated causal genes, and can complement Genome-Wide Association “discovery” approaches, which so far have yielded highly polygenic signals elsewhere in the genome that explain part of the observed phenotypic variation, but are not easily connected to specific effects on phenotypes or functions (Delgado et al., 2011; Dittberner et al., 2018). We can start accounting for this polygenic complexity by integrating genetic information and a functional understanding of the entire genetic developmental pathway, and directly ask if and how known stomatal development genes may promote adaptation.

Here, we use historical specimens as “witnesses” of the ongoing climate change together with molecular genetics knowledge to ask: Can herbaria reveal climate change adaptation in the genetic pathways of essential plant features such as stomata? Can combination of historical and modern genomes with genes’ known phenotypic effects circumvent currently lacking historical phenotyping to predict plant change?

## Results and Discussion

### Strong purifying natural selection defines stomata genes

To investigate how stomata, or stomatal development, has responded to climatic change, we created a novel temporal dataset (see (Latorre, Lang, and Burbano 2022; Lopez et al. 2022)) of 191 globally distributed historical samples, covering the time period from 1817 to 2010 (**Fig. S1A**, **Table S1**), and paired it with the contemporary *A. thaliana’s* 1001 genomes resource (1001 Genomes Consortium 2016). So-called ancient DNA, retrieved from the historical herbarium specimens, was authenticated following the field’s standards (e.g. (Latorre et al. 2020)). Samples show the expected patterns of age-related DNA fragmentation (merged fragments’ median size 98 bp, **Fig. S1B,C**) and damage, i.e. cytosine deamination (reflected as C-to-T substitutions in sequencing data; 0.7-4%, **Fig. S1D**), accumulated particularly at DNA molecules’ termini. As previously shown, deamination of a fragment’s first base is highly correlated with the year of sample collection (one-sided Pearson’s correlation test, correlation coefficient r = −0.589, p = 1.693 × 10^−15^, **Fig. S1E**), a pattern that reflects post-collection aging of the specimens as a primary cause of DNA damage, but whose strength depends on multiple additional factors such as plant specimen collection and storage conditions (Weiß et al. 2016; Sawyer et al. 2012).

Because stomatal density differences have been observed across geographic regions of *A. thaliana* and these changes are partially heritable (Delgado et al., 2011; Dittberner et al., 2018), we aimed to depict genetic variation in known genes of the stomatal pathway, which could constitutively alter stomatal densities. We surveyed the literature and defined a set of 43 stomatal genes (see **Table S2**) that focuses on genes mediating the cell divisions and fate transitions central to stomatal development (**Fig. 1a**). Aiming to specifically inquire if the central developmental pathway itself creates constitutive stomata density differences that can undergo positive selection, we excluded genes that are predominantly characterized as environmental sensors affecting stomatal stress responses.

**Fig. 1:**
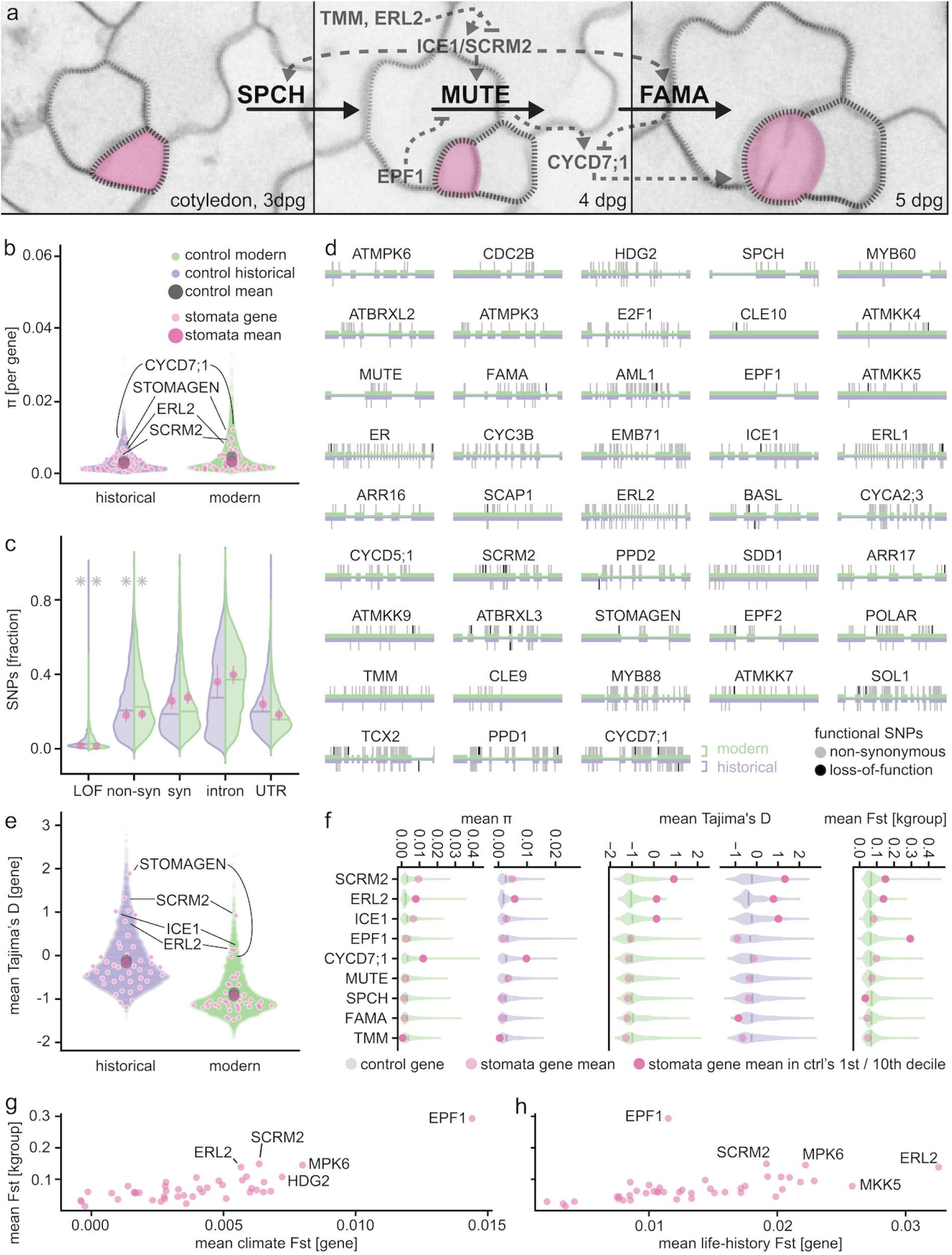
Core stomata genes are conserved while regulatory genes show signals of local adaptation. (**a**) Stomata in A. thaliana are generated through a set of defined symmetric and asymmetric cell divisions that are orchestrated by a well-studied gene network (simplified signaling cascades, with example regulators in gray, selected based on results shown in Fig. 1F, and central (core) transcription factors SPCH, MUTE and FAMA in black. This study focuses on 43 genes central to stomatal development selected based on molecular and experimental studies (**Table S2**). Developmental lineage of a stomatal complex false-colored in magenta. (**b**) Genetic diversity in stomatal genes (single genes light, stomatal gene mean dark pink) is significantly lower than in length-matched control genes (purple for historical, green for modern dataset and gray for dataset mean; outliers marked by gene names; nucleotide diversity π per gene, p_mod_ = 0.004, p_hist_ = 0.046). Asterisks denote significant differences between core and control genes. (**c**) Of the nucleotide polymorphisms present in stomatal genes, significantly fewer are assigned as putative loss-of-function or non-synonymous, and significantly more as synonymous, intronic or located in UTRs than in the control gene distribution (p_mod_^non-syn^ = 0.011, p_hist_^n¤n-syn^ = 0.003, p_mod_^LOF^ = 0.003, p_hist_^LOF^ = 0.049, see **Table S3**). (**d**) Location of LOF (black) and non-synonymous (gray) SNPs in the 43 focus genes in historical (bottom, purple) and modern (top, green) dataset. (**e**) Mean per-gene Tajima’s D as indicator of selection signals, in pink for stomatal genes (and gene group mean in dark pink), purple and green for historical and modern length-matched control-gene distributions. Stomatal genes as a group are not significantly different (p_mod_, p_hist_ > 0.1) from the control, but several genes are outliers in the distribution (marked by gene names). All significance tests for panels **b, c, e** asked if means of the group of stomatal genes were outliers compared to the means of 1000 control groups of 43 length-matched genes each. (**f**) Mean per-gene values for nucleotide diversity π, Tajima’s D and F_ST_^kgroup^ values for outlier genes, in comparison with conserved core stomatal development factors SPCH, MUTE and FAMA (values for all 43 genes, see **Fig. S2**, **S3** and **Table S2**). F_ST_^kgroup^ is calculated for populations defined by whole-genome genetic variation (from (Exposito-Alonso, Vasseur, et al. 2018)). Gene values are displayed as transparent pink circles on top of violin plots representing distribution of values for the respective length-matched control genes. Solid pink circles indicate that the gene mean value lies within the 1st / 10th decile of the control distribution. (**g**) Stomatal gene differentiation as measured by mean F_ST_ per gene, with F_ST_^kgroup^ (y-axis), compared with F_ST_for populations clustered by climate of origin (precipitation and temperature as defined in BIO4 and BIO15, Bioclim dataset, (Hijmans, Cameron, and Parra 2005)), and (**h**) life-history traits (life-history data: (Exposito-Alonso 2020)). Genes with the highest F_ST_-values across the three analyses are labeled.

We hypothesized that if there was an increased genetic diversity in these 43 stomatal genes, it might reflect broad variation in stomatal size and density (the number of stomata per plant surface area) that may have allowed local adaptation to the different environments encountered by the species across its geographic range. To investigate this, we examined our modern and historical genomes as “snapshots” of the species’ current and past ~200 years of global genetic diversity (**Fig. 1b, c**). Overall, expectedly, we found fewer single nucleotide variants (SNPs) in the 191 historical than the 1,135 modern samples. This is consistent with the historical dataset’s much smaller size and stringent SNP-calling that has to account for characteristics and sequencing challenges intrinsic to historical DNA (**Fig. S1B-D**; e.g. (Latorre et al. 2020)). In addition, a modern increase in SNP numbers is also consistent with a recent human-linked population expansion of the species (C.-R. Lee et al. 2017; Exposito-Alonso, Becker, et al. 2018). To compare genetic diversity in these genes in the context of the broader genome, we generated 1000 sets of 43 randomly drawn control genes, with a gene length distribution matched to that of the 43 stomatal genes. In comparison to this control, stomatal genes harbor drastically reduced genetic diversity, as estimated by Watterson’s θ (**Fig. S2**; (Watterson 1975); p_mod_ = 0.006, p_hist_ = 0.069), and lower nucleotide diversity π (**Fig. 1b**, **S2**; (Hahn 2018), p_mod_ = 0.004, p_hist_ = 0.046). This would be expected if purifying selection is purging genetic variation in the developmentally important stomatal genes.

Analyses of non-synonymous and synonymous SNPs in stomata and control genes further show that there are significantly fewer variants annotated as potentially affecting protein function, which we expect to be generally detrimental (partial or full loss-of-function from missense, frameshift, or gain of stop codon), compared to likely neutral variants (located in introns, UTRs, or degenerated codons; annotation with SnpEff (Cingolani et al. 2012); p_mod_^non-syn^ = 0.011, p_hist_^non-syn^ = 0.003, p_mod_^LOF^ = 0.003, p_hist_^LOF^ = 0.049; **Fig. 1c**, **Table S3**). Despite the overall low variation, in the modern data all but one stomatal genes do harbor non-synonymous variation, and some even LOF variants at low frequency (non-synonymous: from 4 (*CLE10*, AT1G69320) to 124 (*CYCD7;1*, AT5G02110), LOF: from 1 to 4, in 23 out of 43 genes; **Fig. 1d, Table S2**). We hereafter refer to non-synonymous and LOF variants jointly as putatively functional variation. As expected, the transcription factors that are essential for stomatal development are among the genes with the lowest genetic variation (SPCH = 0.006 SNPs/bp, MUTE = 0.008 SNPs/bp, FAMA = 0.008 SNPs/bp; **Fig. 1d,** genes ordered by modern data-based SNPs/bp, from left to right, top to bottom). We concluded that stomata genes are generally under purifying selection – especially master regulators that lack variation and are unlikely involved in local adaptation – although some genes still harbor non-synonymous variants at low frequency that could have strong phenotypic effects.

This led us to think of the stomatal pathway as composed of two main groups of genes. One group represents highly conserved essential core genes, where the functional loss of any single gene drastically affects stomatal morphology and density, or is even lethal (MacAlister, Ohashi-Ito, and Bergmann 2007; Pillitteri et al. 2007; Ohashi-Ito and Bergmann 2006). Here, we consider the above-mentioned non-redundant master transcription factors SPCH (AT5G53210), MUTE (AT5G53210) and FAMA (AT3G24140). The second group consists of their direct and indirect regulators that fine-tune the core pathway’s activity and outcome, including duplicated, redundant genes in the pathway that as pairs – but not individually – are similarly essential (**Fig. 1a**). Since loss or functional changes of single “regulator” genes have on average less impact on overall plant development, they are under less evolutionary constraint and thus may be more likely to harbor genetic variation available for positive selection.

### Local and temporal adaptation signals in stomata regulator genes

If any of the functional natural variation in stomatal genes is adaptive, population genetics statistics might detect potential signals of selection and population differentiation. We used a battery of Tajima’s D and F_st_ statistics among populations grouped by population history, phenotypes, or climates. Tajima’s D, which aims to distinguish between neutrally evolving polymorphic sites and those that may be under positive or balancing selection (Tajima 1989), does not significantly differ between the groups of stomatal and control genes (**Fig. 1e**, p_mod_, p_hist_ > 0.1). However, when comparing each individual stomatal gene with its specific control distribution, we saw a negative Tajima’s D value for the essential core transcription factors SPCH, MUTE and FAMA (**Fig. 1f**, **S2**, **Table S2**). This fits with their key roles as necessary and sufficient drivers of stomatal development (MacAlister, Ohashi-Ito, and Bergmann 2007; Pillitteri et al. 2007; Bergmann, Lukowitz, and Somerville 2004; Ohashi-Ito and Bergmann 2006). In contrast, we identified several outliers among the regulatory genes with both high π, Tajima’s D, and population differentiation measured by Wright’s F_ST_ between geographically separated *A. thaliana* populations (F_ST_^kgroup^, based on k = 11 groups, (Exposito-Alonso, Vasseur, et al. 2018)). The combination of high species-wide Tajima’s D and high cross-population F_ST_ may reflect above-average differentiation of alleles between populations relative to within-population differentiation, and maintenance of multiple alleles of the same gene that may be involved in local adaptation. We find indications of this in several genes, all among the 10% highest values of the control distribution for either one or both Tajima’s D and F_ST_^kgroup^ (**Fig. 1f**, **S2**): *EPF1* (AT2G20875), *ERL2* (AT5G07180) – which also has the second highest π after *CYCD7;1, ICE1* (AT3G26744), *SCRM2* (AT1G12860) and *STOMAGEN/EPFL9* (AT4G12970).

If these outlier genes are indeed involved in local adaptation, the distribution of their genetic variation may follow environmental (climate) gradients such as variability in temperature and humidity (Dittberner et al. 2018; Yan, Zhong, and Shangguan 2017). We tested this by grouping samples into populations either based on their collection locations’ precipitation and temperature seasonality (BIOCLIM 4 and BIOCLIM 15, (Hijmans, Cameron, and Parra 2005)), or based on plant life history traits typically aligned with climate, such as the timing of germination or flowering for optimal survival and reproductive success in a given environment (**Fig. 1g, h**, **S3**; (Exposito-Alonso 2020)). Using F_ST_ statistics, we then assessed genetic differentiation between the resulting populations. Despite variation in the absolute F_ST_ values, there is substantial overlap in the genes that are most differentiated between populations, independent of using ancestry, climate or life-history to delineate said populations, with *SCRM2* and *EPF1* among the top four climate differentiators (**Fig. 1g**, **S3**), and *SCRM2* and *ERL2* in the top four life-history genes (**Fig. 1h**, **S3**; see also **Text S1**, **Fig. S6b,c,d** for stomatal gene differentiation over time). Single loss-of-function of the top-four adaptation candidates *SCRM2, ICE1, ERL2, EPF1* tends to minimally affect plant development beyond the stomatal context (Shpak et al. 2005; Hara et al. 2007; Kanaoka et al. 2008), with the exception of *ICEΓs* role in endosperm breakdown and embryo development (Denay et al. 2014). Interestingly, the bHLH transcription factors and paralogs *ICE1 (SCRM*) and *SCRM2*, both involved in cold tolerance, act redundantly as direct interaction partners and expression regulators of *SPCH, MUTE* and *FAMA* (Kanaoka et al. 2008). This fits a model of strong purifying selection in indispensable master transcription factors, and local adaptation through contribution of flexible regulators.

### Functional polygenic scores recover phenotype changes over geographic space

Although the stomatal development pathway at first glance may appear relatively simple, leaf stomatal density is a complex trait. It is likely affected by a combination of effects and complex fine-tuning of the aforementioned core and regulatory genes and likely many others. Genome-Wide Associations of stomatal density thus far yielded moderate heritabilities across many genomic regions, explaining only small fractions of the variation with high uncertainty on specific causal variants (Dittberner et al. 2018). As we have causal evidence that well-known stomatal genes affect the trait, we study nucleotide variants (SNPs) located in coding regions (CDS) where we infer strong protein amino-acid composition change, i.e. non-synonymous changes leading to loss- or gain-of-function. If such putatively functional SNPs are then under climate-driven natural selection, we should expect them to follow geographic and historical climate gradients, and that the non-reference functional variants in multiple genes appear in concerted fashion (Berg and Coop 2014). Indeed, when visualizing the distribution of such variants in six genes with the most putatively functional SNPs, they appear geographically segregated (**Fig. 2a**).

**Fig. 2:**
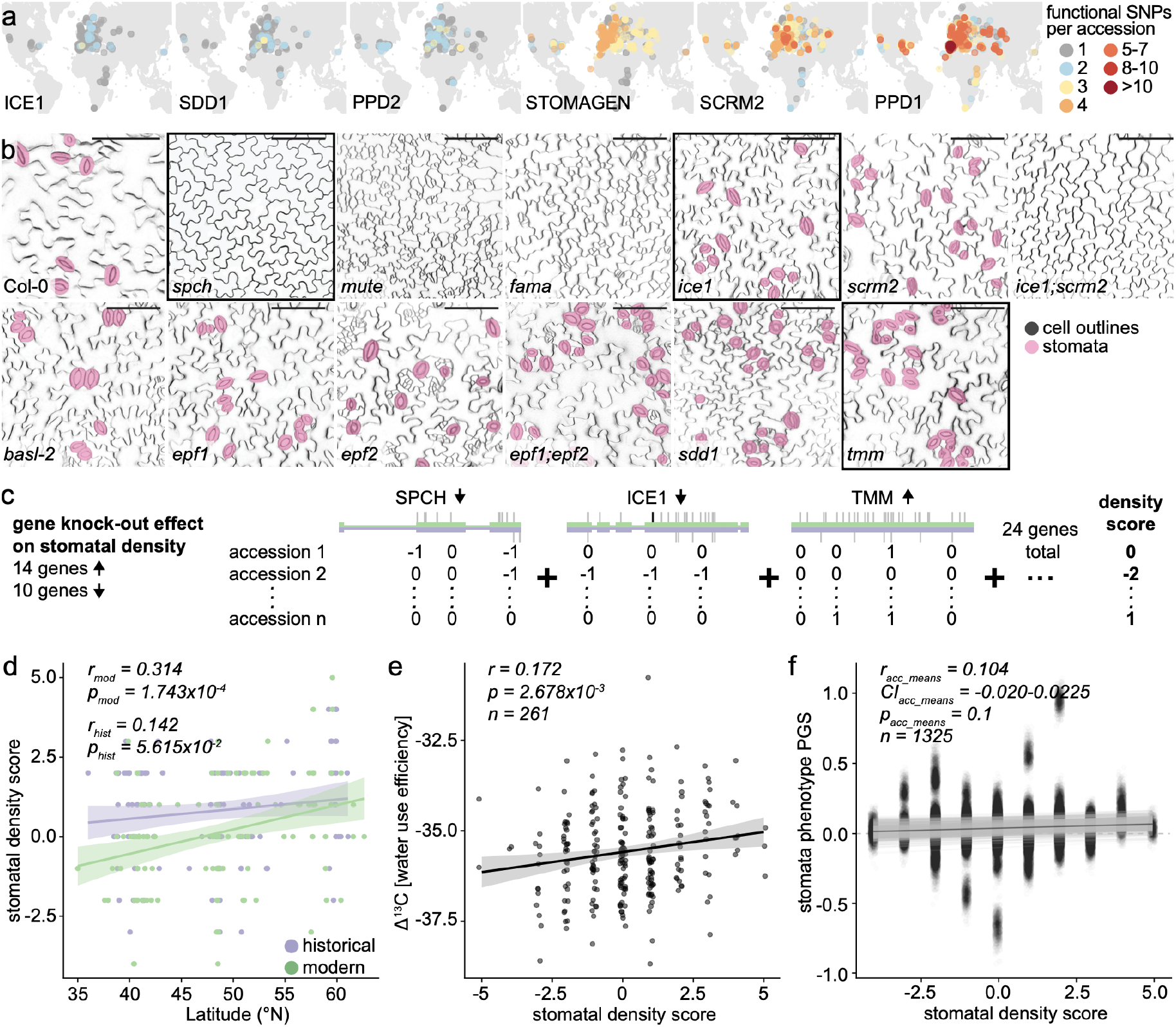
Experimentally-informed polygenic score predicts stomatal density patterns. (**a**) Geographic distribution of functional (non-synonymous, LOF) SNP accumulation in historical and modern samples for six genes with the overall highest amounts of putatively functional SNPs, overlaid on a continental map. Color gradient from gray to dark red indicates samples with 1 to over 10 functional SNPs. (**b**) Gradient of stomatal density differences resulting from loss of major stomatal development genes visualized with confocal microscopy. Stomata are false-colored in magenta, scale bar represents 100 μm. Black frames around three example genes used in **c**. (**c**) Schematic overview detailing the generation of the experimentally informed stomatal density score. Of 24 genes with known effect on stomatal density, loss of 14 increases and loss of 10 decreases stomatal density. Putatively functional SNPs are assigned a ‘-1’ when located in density-decreasing genes and a ‘+1’ in density-increasing genes. Density scores for each historical and modern sample are calculated as the sum of these values across the 24 genes, counting a single functional SNP per gene. (**d**) Linear regression of the stomatal density score with paired samples’ latitude of origin, separated into historical (purple) and modern (green) samples (one-sided Pearson’s correlation test p_mod_ = 1.743 10/ p_hist_ = 5.615 10^−2^, correlation coefficient r_mod_ = 0.314, r_hist_ = 0.142). Analyses exclude samples from North America and the African continent. (**e**) Correlation of the density score with Δ^13^C measured in 261 accessions (one-sided Pearson’s correlation test p = 2.678 10^3^, correlation coefficient r = 0.172; data from (Dittberner et al. 2018)). (**f**) Correlation of the stomatal density score with Genome-Wide Association-based traditional PGS (one-sided Pearson’s correlation test, positive correlation 997/1000 re-trainings, 141/1000 significant one-sided Pearson’s correlation tests with p<0.05, correlation coefficient r_median_ = 0.092).

In our set of 43 stomatal development genes, 24 have confirmed phenotypes in increasing or decreasing stomatal density (mostly shown by knock-out mutant analysis, see exemplary phenotypes in **Fig. 2b**, (Yang and Sack 1995; Berger and Altmann 2000; Nadeau and Sack 2002; Bergmann, Lukowitz, and Somerville 2004; Boudolf et al. 2004; Ohashi-Ito and Bergmann 2006; MacAlister, Ohashi-Ito, and Bergmann 2007; Pillitteri et al. 2007; Kanaoka et al. 2008; Hara et al. 2009; Sugano et al. 2010; Lampard, Wengier, and Bergmann 2014; Gonzalez et al. 2015; Castorina et al. 2016; Han et al. 2018; Vatén et al. 2018; Zoulias et al. 2018; Rowe et al. 2019)). We combined this information on functional variation in the 24 stomatal density genes in our historical and modern samples with the genes’ putative effects on increasing or decreasing stomatal density. With this genetic and phenotypic information, we devised a simple cumulative “stomatal density score”, which is conceptually similar to polygenic scores but is based on summing over putatively functional variation of genes with known increase/decrease effects on a trait. Putatively functional SNPs are assigned a ‘−1’ in genes whose knock-out decreases stomatal density (10 genes), and a ‘+1’ in genes whose knock-out increases stomatal density (14 genes, **Table S4**), where genes with multiple SNPs are only counted once (**Fig. 2c**). This recovered a gradient of “stomatal density scores” (**Fig. 2d-f**).

We found that this density score correlates with samples’ latitude of origin, which in turn dictates life cycle length (**Fig. 2d**, one-sided Pearson’s correlation test p_mod_ = 1.743 10^−4^, p_hist_ = 5.615 10^−2^, correlation coefficient r_mod_ = 0.314, r_hist_ = 0.142; correlation consistently significant upon population structure correction, see also **Text S2**). In fact, experimentally-measured stomatal density also follows a latitudinal trend attributed to *A. thaliana’s* ecological and life-history adaptations to latitudinal climate gradients (**Fig. 2d, Fig. S4a,b**; (Dittberner et al. 2018; Debieu et al. 2013; Exposito-Alonso 2020)). Despite the score being derived from simple functional single-gene knock-out experiments, we found it correlates with a measure of integrated life-time water use efficiency based on CO_2_exchange and water loss (**Fig. 2e**, Δ^13^C, the ratio of ^13^C to ^12^C; one-sided Pearson’s correlation test p = 2.678 10^−3^, correlation coefficient r = 0.172). A positive trend was found with directly measured stomatal density (**Fig. S4c**; Δ^13^C and stomatal density data from (Dittberner et al. 2018)). Permuting the functional (positive or negative) effects of the 24 genes and recomputing the density score removed all significant relationships above (**Fig. S4d,e**), indicating that the density score is not a result of internal dataset biases or population structure. Finally, we compared our functionally-derived density score with a traditional GWA-based polygenic score (PGS; (Chang et al. 2015; Choi, Mak, and O’Reilly 2020)) based on published stomatal density data (Dittberner et al. 2018). PGS trained on 80% of the data explain on average 9.2% of the variance in the reserved 20% of phenotype data (positive correlation in 1000/1000 re-trainings, 745/1000 significant one-sided Pearson’s correlation tests with p<0.05, correlation coefficient r_min-max_ = 0.010−0.561, r_median_ = 0.304; **Fig. S5a,b**). They also correlated with the functionally-derived density score (on average 0.9% of the density score’s variance explained; positive correlation in 997/1000 re-trainings, 141/1000 significant one-sided Pearson’s correlation tests, p<0.05, correlation coefficient r_min-max_ = −0.020−0.204, r_median_ = 0.092; **Fig. 2f, Fig. S5c**; correlations with experimentally measured density and density score lost upon randomization of the phenotype-genotype associations in the training dataset, **Fig. S5d,e**, **Table S5**).

### Polygenic score predicts historical reduction of stomatal density

Ultimately, we aim to understand how stomatal variation in *A. thaliana* may have changed over the last centuries of climate change. For this, we calculated our stomatal density score on historical genomes. To avoid geographical biases past and present, we made comparisons within a subset of historical and modern samples, paired based on geographic proximity (126 pairs, minimum-maximum distance = 0.5-495 km, mean = 139 km; Pearson’s correlation of sample pairs’ latitudes of origin, r = 0.990, p < 2.2 10^−16^, **Fig. 3a, Fig. S6a**, **Table S6**). The historical set for these comparisons ranged between 1817 and 2002 (mean collection year 1923), and the modern set between 1992 and 2012 (mean 2001), with a mean age difference of 79 years in historical-modern sample pairs (from 1 to 182 years).

**Fig. 3:**
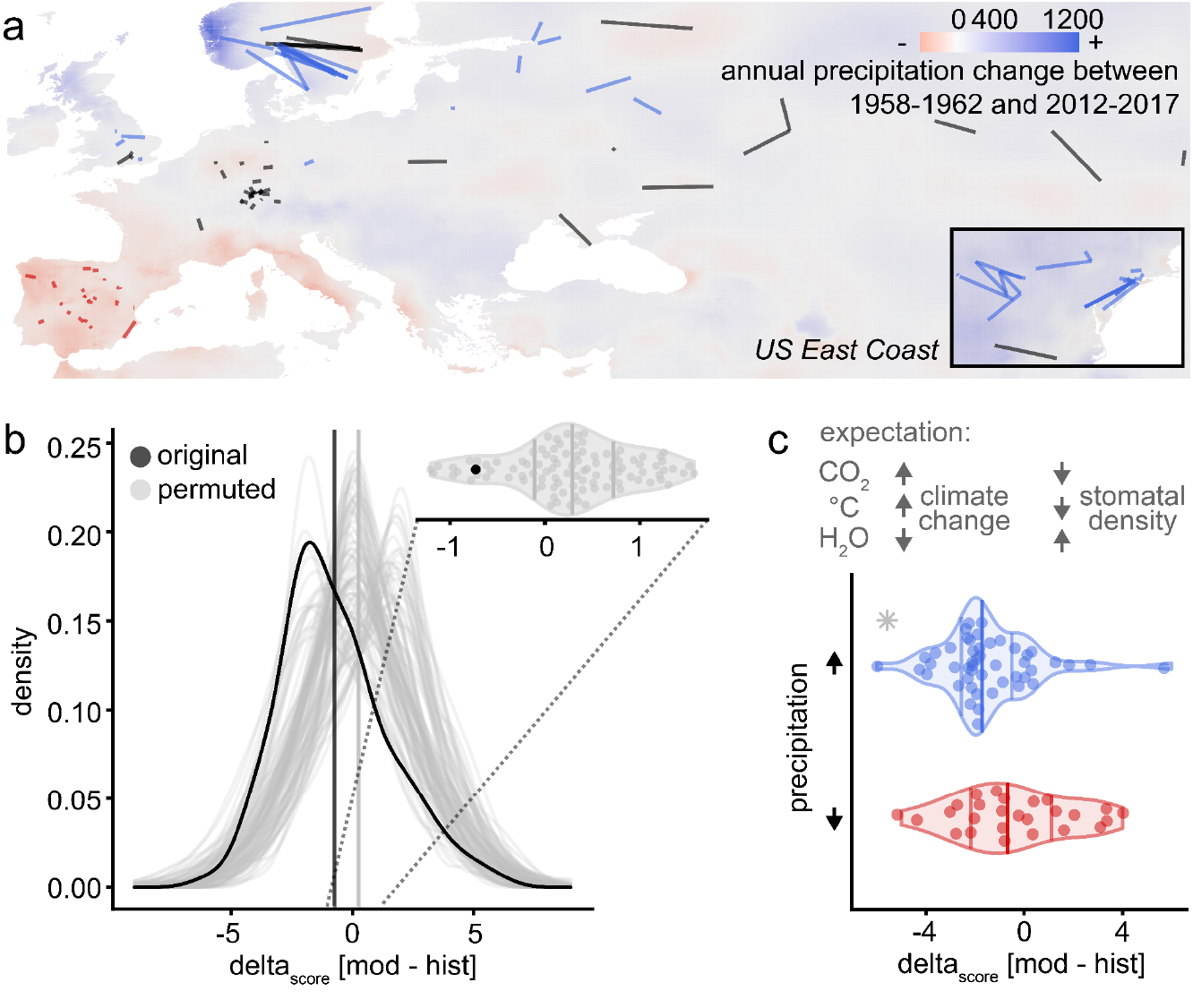
Stomatal density decrease over time fits climate change expectations. (**a**) Map with historical and modern sample pairs as used for stomatal density change analyses (see **b**, **c**). Connecting lines between sample pairs are colored by the precipitation change in the sample locations, red indicates a significant decrease in precipitation from 1958 to 2017 in both the historical and modern sample location, blue indicates a significant increase in precipitation, and black no significant change or different changes in paired locations. Background color gradient indicates change in the mean annual precipitation between 1958-1962 and 2012-2017, with colors as above. Inlay shows sample pairs located at the Northern American East Coast. (**b**) Distribution of per sample-pair calculated difference in stomatal density scores (delta_score_) for original data (black) and 100 permutations (gray) of genes and their assigned effect (decrease/increase) on stomatal density; mean delta_score_ = −0.707, Wilcoxon signed rank test, p < 2.2 10^−16^. Density distribution means are marked by solid black and gray vertical lines. Violin plot of the distribution means, with the non-permuted mean delta_score_ lower than 92/100 permutations. (**c**) Expected effects of climate-change related shifts in CO_2_ concentration, temperature and water availability on stomatal density (based on published experiments, e.g. (Crawford et al. 2012; Vile et al. 2012; Yan, Zhong, and Shangguan 2017; Lau et al. 2018)). Change in the stomatal density score (deltascore) in sample pairs with significantly increased (blue) or decreased (red) precipitation in geographic locations of origin (excluding sample pairs where precipitation did not change significantly, or where a pairs’ locations did not change in the same way). Increased, but not decreased precipitation is significantly associated with decreased delta_score_ (linear regression, delta_score_ ~ precipitation_directionality_, p-value_incrprecipitation_ = 0.024, p-value_decr___precipitation_ = 0.889, p-value_mode_? = 0.045; see also **Table S9**). Analyses include samples from North America and exclude samples from the African continent as well as pairs between island and mainland samples.

Before calculating temporal trends, we established that the stomatal density score correlation with latitude remains consistent also in the historical dataset, suggesting patterns of local adaptation have long been established in the species (see historical trends in **Fig. S4a,b,d,e**). We then calculated the difference in density score (delta_score_) for each individual historical-modern geographic sample pair (**Fig. 3a**). The resulting distribution of score changes corroborates prior results that suggest a decrease of stomatal density in modern samples (black distribution, mean delta_score_ = −0.730, Wilcoxon signed rank test, p < 2.2 10^−16^, **Fig. 3b**, **S7d**; this decrease was robust to removal of samples along a latitudinal gradient, or modern samples with uncertain collection dates, mean delta_score_^noUS^ = −0.433, delta_score_^noUS^-^noSpain^ = −0.423, delta_score_^noScandinavia^ = −0.745, delta_score_^noUncertainty^ = −0.717, Wilcoxon signed rank test all p < 1.25 10^−15^, **Fig. S7a-c**, **Table S7, S8**, see also **Text S3**). Conducting again permutations of positive and negative stomatal density gene effects confirmed that the decrease in stomatal density is only recovered with coordinated loss or gain of certain functional variants, rather than being based on confounders such as whole-genome divergences (**Fig. 3b**, **S7a-d**, gray distributions).

A decrease in stomatal density over the last century fits the expectation based on experimental evidence showing increases in CO_2_ concentration and temperatures leading to leaves with lower stomatal density (Woodward 1987; Van Der Burgh et al. 1993; McElwain and Chaloner 1995; Y. Li et al. 2014; Samakovli et al. 2020; Lau et al. 2018; Crawford et al. 2012). However, while CO_2_ concentration and temperature have mostly increased, climate change has also altered weather patterns and heterogeneously increased or decreased precipitation depending on the region on Earth (Pörtner et al., n.d.). Because experiments simulating decreased water availability show directly opposite stomatal responses (density increases, e.g. (Doheny-Adams et al. 2012; Vile et al. 2012)), we wondered whether we could use the differential changes in precipitation as a natural counterfactual experiment to validate our genetic prediction of stomata changes over time. We thus conducted a sensitivity analysis, grouping sample pairs based on their temperature and precipitation trajectories and magnitudes of change over the last 60 years (**Fig. 3a,c**). Sample pairs from locations with increased precipitation are significantly more likely to have decreased stomatal density score over time (**Fig. 3c**, **Table S9**; **Fig. S7e**, 2.5 fold odds of stomatal decrease with Fisher’s Exact Test, odds_ppt-high_ = 2.415, p_ppt-high_ = 0.025, and consistent results for the same analysis with less stringent filters in **Fig. S7f (Abatzoglou et al. 2018)**; **Table S10**; see also **Text S3**). Locations with decreased precipitation, counteracting the expected effects of increased CO_2_ and temperature, displayed a (non-significant) trend of stomatal density increase (Fisher’s Exact Test, odds_ppt_low_ = 0.589, p_ppt_low_ = 0.283, **Table S10**; e.g. (Doheny-Adams et al. 2012; Vile et al. 2012)).

Despite the spatial sample pairing and climate splits in our analyses, temporal trends in stomatal density score changes might still contain residual biases such as population structure. We therefore conducted a series of analyses including fitting the first three main genomic principal components (PC) to capture population structure, and describe the stability of various estimates (**Text S3**). The signal of lowering average stomatal density remains after population structure correction (p-value_model2_ = 0.001, **Text S3, Table S9**), and also the more pronounced decrease in stomatal density in regions with increased precipitation is mostly consistent after corrections for each PC axis (p-value = 0.034, 0.081, 0.019, after PC1, PC2, PC3 corrections, while no variable is significant with a full model; **Text S3, Table S9**). Taken together, while the signal strength is variable, we consistently identify trends of decreasing stomatal density that correspond well both with experimental and historical observations of stomatal density responses to the climate variation connected with global change.

### Conclusions and outlook

Global change has led to rapid and drastic changes of multiple climate parameters –CO_2_ concentration, temperature, precipitation –that have a strong impact on plant development. Here, we studied responses of *A. thaliana* leaf stomatal development to anthropogenic climate change, using historical herbarium genomes as witnesses of this multi-factorial “global change experiment” (e.g. (Woodward 1987; DeLeo et al. 2019)). Despite the overall high conservation of stomatal development genes, our integrative approach allowed us to identify several genes with evolutionary signals consistent with local adaptation across *A. thaliana’s* geographic range, suggesting stomatal development could evolve to climate change conditions. We developed a novel polygenic score based on functional molecular knowledge of stomatal development genes that agrees with a historical trend of stomatal density decrease in *A. thaliana*, classically observed across species but of unknown genetic basis (Woodward 1987; Beerling and Chaloner 1993). While evolutionary processes can be fast (e.g. (Exposito-Alonso, Becker, et al. 2018; Rudman et al. 2022; Franks et al. 2016)), our analyses show that even with hundreds of historical genomes, the described trends of putative adaptive evolution are close to the detection limit above the noise of genetic drift, phenotypic plasticity, and methodological idiosyncrasies, as shown by the re-analyses with different geographical subsets of the data. Ultimately, our discovery will stimulate follow-up investigations such as studying the molecular mechanisms of how these genes promote adaptation –now enabled for instance by single base-editing with CRISPR to recreate historical variants (Tan et al. 2020; Kang et al. 2018) -, characterizing cellular phenotypes directly from historical specimen tissues using customized microscopy techniques (e.g. (Haus, Kelsch, and Jacobs 2015)), and generating denser timelines of historical genomes possible with high-throughput aDNA technologies (Gutaker and Burbano 2017; Kistler et al. 2020). The functional historical genomics approach presented here adds a new exciting avenue to leverage the power of genomics to reconstruct phenotypic impacts of climate change on species, even for those phenotypes that are cellular or sub-cellular. Ultimately, our approach could help uncover complex responses involving water-use efficiency, photosynthetic capacity or drought-resistance that are not preserved or measurable in historical collections. Disentangling this will be key to understanding plants’ past and future adaptive potential and design targets for engineering plants for the future.

## Materials and Methods

### Sequence data

#### Contemporary data

Published contemporary, SnpEff-annotated (Cingolani et al. 2012) *Arabidopsis thaliana* single nucleotide variants (SNPs, (1001 Genomes Consortium 2016)) were annotated (gene, CDS, etc.) with the publicly available TAIR10 release (TAIR10_GFF3_genes_transposons.gff, link, (Berardini et al. 2015)). The as “gene” annotated fraction of SNPs was extracted for subsequent analysis using bash, BCFtools v1.10.2 (Danecek et al. 2021), PLINK v1.90b6.16 64-bit (Purcell et al. 2007; Chang et al. 2015) and R (R Core Team (2019). R: A language and environment for statistical computing. R Foundation for Statistical Computing, Vienna, Austria. https://www.R-project.org/) within RStudio v1.2.1335 (RStudio Team (2018). RStudio: Integrated Development for R. RStudio, Inc., Boston, MA. http://www.rstudio.com/). Samples lacking coordinates for their geographic origin or information on the year of their original collection were excluded for some analyses as indicated.

#### Herbarium data

##### Sample processing and sequencing

Whole-genome sequenced African (n = 9), North American (n = 33), German (n = 34) and global (n = 204) *A. thaliana* herbarium specimens were downloaded from public repositories (Exposito-Alonso, Becker, et al. 2018; Durvasula et al. 2017; Latorre, Lang, and Burbano 2022; Lopez et al. 2022).

All historical sequencing data was processed as described (Latorre et al. 2020; Meyer and Kircher 2010). Briefly, after trimming and merging (Adapterremoval v2.3.1, (Schubert, Lindgreen, and Orlando 2016)) the demultiplexed reads, we mapped the merged read fraction (BWA v0.7.15-r1140, (H. Li 2013)) to TAIR10 (Berardini et al. 2015), filtered for mapping quality >=20 (Samtools v1.9, (Danecek et al. 2021)), removed PCR duplicates (DeDup v0.12.8, (Peltzer et al. 2016)) and confirmed the authenticity of our historical samples by establishing fragmentation (Samtools v1.9, (Danecek et al. 2021); median merged fragment length 98 bp, insert-sizes for sheared, unmerged fragments 234 bp) and deamination damage patterns (MapDamage v2.2.1, (Jónsson et al. 2013); **Fig. S1**). For double-processed (sheared and unsheared) global samples (Lopez et al., unpublished), we use Samtools to concatenate the merged fraction of unsheared samples together with both the merged and unmerged fraction of sheared samples into a single file.

Based on sequencing and mapping statistics obtained through Samtools (stats), DeDup and MapDamage, we defined quality thresholds. We retained only samples with >1,000,000 bp sequenced, >10,000 reads sequenced, >95% of reads mapped, a duplication rate <0.3, an error rate <0.02 for *de novo* SNPcalling, and excluded samples with >0.5 missing genotypes, resulting in a total of 191 samples. We did not differentiate between “regular” (Lopez et al. 2022; Latorre, Lang, and Burbano 2022) and UDG-treated libraries (Exposito-Alonso, Becker, et al. 2018; Durvasula et al. 2017). Samples lacking coordinates for their geographic origin or information on the year of their original collection were excluded for some analyses as indicated.

##### De novo SNPcall

For genome-wide *de novo* calling of SNPs in the full historical dataset, we created and indexed a reference dictionary with TAIR10 (Berardini et al. 2015) using Picard’s CreateSequenceDictionary (Picard v2.18.29-0; “Picard Toolkit.” 2019. Broad Institute, GitHub Repository. https://broadinstitute.github.io/picard/; Broad Institute) and samtools faidx (Samtools v1.9, (Danecek et al. 2021)). We then called haplotypes *de novo* (i.e. not based on an existing set of previously called SNPs) using the GATK HaplotypeCaller, and combined resulting .gvcf-files into a single input file with GenomicDBImport before calling variants with GenotypeGVCF, following the *best practice* recommendations for GATK (gatk4-4.2.0.0.-0, (Van der Auwera and O’Connor 2020)). Keeping only single nucleotide variants (*vcftools --remove-indels*, VCFtools/0.1.16, (Danecek et al. 2011)), we then generated two subsets, either including only, or excluding all sites present in the published 1001G vcf-file (*vcftools --positions*; (1001 Genomes Consortium 2016)). To determine parameters for quality-based filtering of the full SNP dataset, we compared quality parameters in the two subsets and the full dataset (*vcftools --gzvcf <infile.vcf.gz> --get-INFO QD --get-INFO FS --get-INFO ReadPosRankSum --get-INFO MQRankSum --get-INFO BaseQRankSum --out <outfile>*) using R and RStudio. Assuming that the distributions of quality parameter values of the 1001-only dataset are representative for high-quality SNPs, we defined cutoff values for quality-based SNP filtering with vcflib (*vcffilter -f ‘DP > 22 & FS < .2 & ReadPosRankSum > (0 –2) & ReadPosRankSum < 2;* vcflib/20161123-git, (Garrison et al. 2021)). Subsequently, we excluded samples with missing call frequencies greater than 50% and filtered for biallelic positions with PLINK (*plink --mind .5;* PLINK v1.90b6.16 64-bit (Purcell et al. 2007; Chang et al. 2015)). Subsequently, we used PLINK to remove variants where less than three individuals carried the alternative allele (*--mac 3*), and samples with site missingness exceeding 15% (*--geno .15*) to generate a more stringently filtered set of high quality SNPs (“hiq-set”). We used variant-based PCAs to visualize the effects of the different filters on the recovered genetic diversity in the samples and identify eventual outliers based on filtering artifacts (*PLINK --pca <sample_number>).* The resulting vcf files were annotated using SnpEff v5.0e (Cingolani et al. 2012) and the TAIR10 *Arabidopsis thaliana* reference genome (Berardini et al. 2015), and further annotated and subset as described above for contemporary data.

#### Joined historical and modern data

Before merging, the historical and modern datasets were individually filtered with PLINK for alleles with a minor allele count of at least 3, and for samples with less than 15% site missingness (see **De novo SNPcall**). Datasets were then intersected using BCFtools (*bcftools isec -n=2*, v1.10.2 (Danecek et al. 2021)) and filtered for the resulting shared SNPs with PLINK (*--extract <shared_SNPs>*) to avoid dataset-specific biases caused by private SNPs. Gene-specific and control subsets were generated as described above, using the same gene lists.

For analyses that directly compare historical and modern samples, we further subset the datasets to only the geographically-matched 126 sample pairs (see **Geographic-distance based sample pairing**) and re-filter for a minor allele count of 3 and lower than 15% site missingness (*PLINK --keep --geno .15--mac 3*).

#### Gene-specific subsets

##### Stomatal gene list

Based on recent literature reviews (Chater et al. 2017; L. R. Lee and Bergmann 2019; Simmons and Bergmann 2016) and lab-internal experience, we generated a list of 43 genes that are central to stomata growth, development and the stomatal lineage (**Table S2**).

##### Control gene lists

Gene-specific features need to be accounted for to generate a subset of control genes that is comparable with the stomata-gene set. The majority of analyses conducted here focuses on statistics on a per-gene level; measurements are thus sensitive to differences between genes in the number of SNPs present, which is strongly correlated with the length of the gene itself. To account for this, control genes are selected on a per-gene basis to match the gene length distribution of the original dataset. We extract gene length information from the gene-subset TAIR gff, thus excluding genome features that have not been annotated as “gene” and making the control dataset more comparable to the stomatal gene set. For one stomatal focus gene at a time, we subsample this gff for all genes of the same length within a margin of +- 2.5%, and of this subset randomly pick one gene, generating a list of 43 control genes, one for each stomatal focus gene. Depending on the analysis, we use 1000 re-samplings of this gene length-based control. The same control-gene sets are used to extract SNPs for all datasets, ie. historical, modern as well as combined dataset, facilitating direct comparisons.

##### Subsetting

For both contemporary and historical data, stomata-specific and control gene lists are used to subset the for “gene” subsampled vcf-files for the specific SNPs, using bash and PLINK v1.90b6.16 64-bit. After annotation of the resulting subsets using R/RStudio, we apply a final filter that excludes any annotations of SNPs in genes not present in our original subset lists. This avoids inflation of (especially control) subsets by SNPs annotated for multiple genes with overlapping genomic locations. Subsequently, we filter annotations for the first and second splice isoforms of the focus genes. For assessing the amount of certain SNP-types (synonymous, non-synonymous, loss-of-function etc.), we count the value of the first splice isoform. Several SNP-types are grouped together to obtain six main classes, all putative based on TAIR and SnpEff annotations: “loss-of-function” (disruptive_inframe-deletion, disruptive_inframe_insertion, inframe_deletion, inframe_insertion, frameshift, start_lost, stop_lost, stop_gained), “non_synonymous” (missense), “synonymous” (synonymous), “UTR” (5_prime_UTR_premature_start_codon_gain, 3_prime_UTR, 5_prime_UTR), “intron” (intron) and “other” (none, splice_region, splice_donor, splice_acceptor, intron, stop_retained, non_coding_transcript_exon, upstream_gene, downstream_gene).

#### Population genetic statistics

All population genetics analyses were analyzed both at a per focus-group and per-gene scale to assess if either one of our stomatal development genes or all genes as a group showed outlier values. When comparing value distributions of stomatal developmental genes (i.e. focus genes) as a group against the random, length-matched control genes (see **Control gene lists**), we calculated mean values for each of the 1000 × 43 control groups and assessed if the stomatal gene group value was higher or lower than the majority of control group means. For per-gene assessments, we calculated mean values for each statistic for each focus gene, and each of 1000 gene-specific length-matched random control genes. To compare focus and control genes, we calculated if the population genetics statistics value of the focus gene is outside of the 0.1 or 0.9 quantile of the control value distribution. All summary calculations, statistical analysis and plotting were done in R/RStudio (R Core Team (2019). R: A language and environment for statistical computing. R Foundation for Statistical Computing, Vienna, Austria. URL https://www.R-project.org/; RStudio Team (2020). RStudio: Integrated Development Environment for R. RStudio, PBC, Boston, MA URL http://www.rstudio.com/).

##### Genetic diversity –π and SNPs per gene

Assessing diversity based on SNPs’ presence, we summed SNPs for each gene and calculated raw SNPs/bp (i.e. the number of SNPs found per gene length) and Watterson’s θ. To estimate nucleotide diversity π per polymorphic site, we used VCFtools default settings (*<--site-pi>;* VCFtools/0.1.16, (Danecek et al. 2011)) and extracted the maximum per-site pi-value per gene. We also calculated 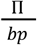 by dividing the sum of all π values for a single gene by the gene length (bp), assuming that all other positions that lack a π-value are invariant with nucleotide diversity 0.

##### Tajima’s D

To identify potential signals of selection in our genes of interest, we calculated Tajima’s D (Tajima 1989) on the full 1001G vcf-file, using VCFtools with the smallest window size of 100 bp (*--TajimaD 100;* VCFtools/0.1.16, (Danecek et al. 2011)). We then associated these values (by definition calculated on a per-window basis) to our biallelic SNPs of interest and summarized them using R and RStudio. We calculated the mean Tajima’s D value per gene for both length-matched control and focus gene sets.

For comparison of Tajima’s D between historical and modern samples we calculated Tajima’s D for the respective independent subsets of samples present in the geographically-paired dataset (see **Joined historical and modern data**).

##### Population fixation index F_ST_

Population differentiation as indicated by the fixation index F_ST_ was estimated with Plink default settings (*plink --fst --within <groups>;* PLINK v1.90b6.16 64-bit (Purcell et al. 2007; Chang et al. 2015)). Population subdivisions for the (“modern”) 1001G dataset were either based on published, genetics-based global *A. thaliana* population structure (11 groups, (Exposito-Alonso, Vasseur, et al. 2018)), on the accessions’ different life-strategies (Exposito-Alonso 2020) or on experienced (micro-)climates (Hijmans, Cameron, and Parra 2005). For the former, admixed individuals as defined in (1001 Genomes Consortium 2016)) were excluded. For the latter, accessions were grouped based on the temperature and precipitation seasonality (see (Dittberner et al. 2018) for environmental conditions correlated with stomatal size/density) of the accession’s latitude/longitude of origin (WorldClim.org) or the accession’s environment-independent germination rate and cold-induced dormancy (as used in (Martínez-Berdeja et al. 2020; Exposito-Alonso 2020), but excluding accessions with imputed phenotypes). To generate clusters for both parameters, we calculated accessions’ silhouette scores (cluster R-library, *silhouette*; Maechler, M., Rousseeuw, P., Struyf, A., Hubert, M., Hornik, K.(2019). cluster: Cluster Analysis Basics and Extensions. R package version 2.1.0.) for two through fifteen clusters and used the number of clusters with the highest score (6 and 4 respectively) in a single k-means clustering. Accessions and their respective cluster identity were combined in a separate fam-format file for the environmental and life history clustering each, and used to calculate F_ST_ as described above for both stomatal genes and 1000 re-samplings of the gene length-based control. All summary calculations, statistical analysis and plotting were done in R/RStudio.

Using F_ST_ to assess the genetic differentiation between populations not across (geographic) space, but over time, we in addition calculated F_ST_ for the paired dataset (see **Joined historical and modern data**), with the samples’ identity as “modern” or “historical” defining the two populations.

#### Matching historical and modern samples

##### Geographic-distance based sample pairing

Historical samples were matched to the geographically closest modern sample from the 1001G *A. thaliana* dataset (1001 Genomes Consortium 2016). For each historical sample, we calculated pairwise distances to the geographic origin of all modern samples with the Vicenty (ellipsoid) method (geosphere R-library, *distVincentyEllipsoid;* Robert J. Hijmans (2019). geosphere: Spherical Trigonometry. R package version 1.5-10. https://CRAN.R-project.org/package=geosphere). We selected the geographically closest modern sample as a “match” and removed this modern sample from subsequent pairwise distance calculations to generate unique historical-modern sample pairs. Historical-modern pairs with a distance larger than 500 km were removed to obtain a final dataset of 126 pairs, filtering out all historical samples from the African continent, as well as pairs between islands and mainland, or matched across bodies of water or mountain ranges (**Table S6**). For some analyses, samples from North America (originating west of longitude -25) were removed as well, retaining 116 sample pairs. Only sample pairs with the historical collection date predating the modern one were considered.

##### Climate change trajectories

Despite this close pairing, sample pairs’ geographic origins may still be sufficiently far apart to have experienced diverging climatic changes in the period of time between historical and modern sampling. Such climatic differences may translate into differing selection pressures and an overall different genetic makeup already in the past, which complicates attributing observed genetic differences between historical and modern samples to temporal (adaptive) processes alone. We reduced these confounding factors by modeling the directionality of climate change between 1958 and 2017 as recorded in the TerraClimate dataset (resolution ~4km, (Abatzoglou et al. 2018)). For the geographic location of each historical and modern sample, we extracted precipitation, and the minimum and maximum monthly temperatures. Temperatures were then subset to one value per year, the month with the highest (tmax) recorded temperature. To extract trends in the monthly precipitation, we transformed records into a time series and decomposed it to separate the gradual trend over time from the periodic seasonal precipitation variation (R/Rstudio, stats-package). With the resulting annual values for tmax and ppt per sample location we assessed directionality (increase or decrease) of an observed climate change by extracting the slope of a linear regression of the climate parameter over time. Using Spearman correlation, we calculated the p-values of these climate change trends, corrected for multiple testing with the Benjamini-Hochberg procedure, and assigned trends that pass a threshold of p_BH_<0.01 as significant. Based on their climate, sample-pairs were then classified as either matching, i.e. showing the same significant increase or decrease in maximum temperature or precipitation, or not showing any significant change, or not matching, i.e. showing opposing directionalities of change, or only one of the two showing a significant change in either direction. This was used to evaluate changes in stomatal density scores over time under specific climate conditions (see **Stomatal density**).

#### Stomatal density

##### Experimentally informed genetics-based stomatal density proxy

To generate a proxy for stomatal density based on genetic information, we searched the literature for the effect a mutation in each of the 43 stomatal genes has on stomatal density: higher density (14 genes), lower density (10 genes), or none (18 genes; see **Table S4**). We then use as protein function affecting (non-synonymous and putative loss-of-function; jointly referred to as putatively “functional”) assigned SNPs (see **Gene specific subsets**) to calculate a stomatal density proxy for each historical and modern accession. Accessions that carry the reference allele for a functional SNP are assigned a value of ‘0’, i.e. no expected effect of this SNP on stomatal density. Presence of the alternative allele translates into ‘+1’ if the SNP is located in a gene whose mutation is known to increase, and ‘-1’ if the gene’s mutation is known to decrease stomatal density. SNPs within the same gene (or in general SNPs in close proximity) may be part of the same haplotype and can thus not be treated as statistically independent. Thus, to calculate a stomatal density score for each sample, we summed the assigned values across all genes with a density phenotype, but took only one functional SNP per gene into account. Scores were calculated for all accessions within the 1001G (1001 Genomes Consortium 2016) and our panel of historical samples.

The stomatal density score was correlated with published experimentally measured stomatal densities and Δ^13^C (Dittberner et al. 2018). Assessment of the score’s correlation with latitude was performed on the full set of 191 historical samples. To account for a potential effect of geographical sampling bias, the modern dataset of 1135 accessions was subsampled 100x without replacement to mirror the size of the historical dataset. Linear regression was calculated for all individual samplesets, and slopes were extracted using R/RStudio. For interpretations of the linear models’ intercepts, we aimed to avoid extrapolation of the association beyond the geographical space covered by samples, and thus re-calculated the intercept using the median latitude of the historical and modern sampleset, respectively. To further assess the contribution of *A. thaliana’s* global population structure to the stomatal density score, we performed 100 permutations of the association of density phenotypes (increase/decrease) and stomatal genes. This was done both to validate the stomatal density score’s latitudinal trajectory, as well as the density shift from historical to modern samples. For the latter, we subtract the historical sample’s score from its modern partner within each geographic distance-based historical-modern sample pair (see **Matching historical and modern samples**). This within-pair calculation aims to reduce effects of population structure differences between historical and modern samples, increasing the probability that identified differences result from actual change over time.

We further subset the sample pairs by the climate change experienced at their geographic location over the last 60 years (see **Climate change trajectories**). Only locations where the change from 1968-2017 is significant are included, to dissect how the stomatal density score is affected by increasing or decreasing precipitation and temperature individually, and in combination. With Fisher’s Exact Test for Count Data (*stats::fisher.test*(), R) we then calculate the odds and 95% CI that stomatal density in modern samples has decreased under the single and combined climate conditions, identifying density score changes for each sample pair individually, and only including sample pairs where both geographic locations have experienced the same significant change in climate over the last 60 years.

##### Traditional polygenic score model

For each of a 1000 iterations, phenotypic data from (Dittberner et al. 2018) was randomly split 4:1 into a training and a test set using the scikit-learn sampling function (Fabian Pedregosa, Gaël Varoquaux, Alexandre Gramfort, Vincent Michel, Bertrand Thirion, Olivier Grisel, Mathieu Blondel, Peter Prettenhofer, Ron Weiss, Vincent Dubourg, Jake Vanderplas, Alexandre Passos, David Cournapeau, Matthieu Brucher, Matthieu Perrot, Édouard Duchesnay 2011). With GEMMA (v0.89.1, (Zhou and Stephens 2012)), we calculated either genome wide associations, or associations with the subset of 43 stomata genes as described above, using a univariate linear mixed model on the training phenotypes and the joined historical and modern genotypic datasets described above.

With Plink (v1.9), we then generated an additive polygenic score (PGS) model on the genome wide/stomata gene associations using a p-score threshold of greater than 0.05 to select the most predictive SNPs. Plink and subsequent R scripts used for PGS analysis followed methods as published (Choi, Mak, and O’Reilly 2020). Phenotypes predicted by the PGS model were then compared to the test set or to the stomatal density score (see **Experimentally informed genetics-based stomatal density proxy**) to determine the accuracy of the model using one-sided Pearson’s correlation tests. In order to test whether the PGS models capture the phenotypes’ genetic signatures, we ran 1000 iterations where the phenotypic data was permuted and reassigned to random genotypes in the training set, and subsequently followed steps for GWA and Plink modeling as described above. For all comparisons from the 1000 iterations, we additionally calculated the mean predicted PGS phenotype per accession and tested these phenotypes’ correlation with measured stomatal density or the stomatal density score with one-sided Pearson’s tests.

##### Plant growth and conditions

*A. thaliana* mutant line seeds were sterilized with chlorine gas (produced by mixing 50 mL bleach with 2.5 mL 37% hydrochloric acid) and stratified on MS plates (½ MS (Caisson Labs), 1% Agar (w/v), pH 5.7) for two days at 4°C before transfer to a 22°C chamber with 16h light : 8h dark cycles (110 μmol m^−2^ s^−1^).

The following mutants and transgenic lines were reported previously: *basl-2* (Dong *et al.* 2009), *epf1-1* (Hara *et al.* 2007), *epf2-1* (Hunt & Gray 2009), *ice1-D (scrm-D*, Kanaoka *et al.* 2008), *ice1-2* (Kanaoka *et al.* 2008), *mute* (Pillitteri *et al.* 2007), *scrm2-1* (Kanaoka *et al.* 2008), *sdd1-1* (Berger & Altmann 2000), *spch-3* (MacAlister *et al.* 2007), *tmm-1* (Yang & Sack 1995), *fama* (Ohashi-Ito & Bergmann 2006), *epf1-1;epf2-1* (Hunt & Gray 2009), and *ice1-2;scrm2-1* (Kanaoka *et al.* 2008). Natural *A. thaliana* accessions were published previously (Col-0, e.g. (1001 Genomes Consortium 2016)).

##### Microscopy and image analysis

To visualize cell outlines in cotyledons, we stained 9 dpg seedlings of *A. thaliana* mutant lines with FM4-64 p (N-(3-triethylammoniumpropyl)-4-(6-(4-(diethylamino)phenyl)hexatrienyl)pyridinium dibromide, ThermoFisher Catalog #T13320). The *spch-3* mutant (MacAlister *et al.* 2007) had been transformed with a plasma membrane marker, *pATML1::mCherry-RCI2A*, and was not stained with FM4-64 prior to imaging. Cotyledons were imaged on a Leica SP5 confocal microscope with HyD detectors using a 40X NA1.1 water objective at a resolution of 1024 × 1024 pixels. Images were post-processed (contrast enhancement and noise reduction) using Fiji (V2.1.0/1.53c; (Schindelin et al. 2012)) and Adobe Illustrator V26.3.1.

## Data availability

*Arabidopsis thaliana* 1001 variant data was published in (1001 Genomes Consortium 2016), and respective kgroups in (Exposito-Alonso, Vasseur, et al. 2018). Historical sequencing data is publicly available (North American accessions: (Exposito-Alonso, Becker, et al. 2018), African accessions: (Durvasula et al. 2017), German accessions: (Latorre, Lang, and Burbano 2022), worldwide accessions: (Lopez et al. 2022)).

## Acknowledgements

We are grateful to members of the Bergmann and Exposito-Alonso labs for support, suggestions, discussion, input for experiments, analysis and reviewing the manuscript, Lucas Czech for bioinformatics and Jeffrey P. Spence for statistics support.

## Supplemental Material

### Supplemental Text

#### Text S1. Temporal differentiation in stomatal genes

We also made use of the paired historical and modern samples to calculate the sample groups’ temporal differentiation, using either F_ST_^tιme^ to measure differentiation between historical and modern populations, or investigating changes in Tajima’s D between the two. *SCRM2, ERL2* and *MYB60*, outliers in one or several of the ancestry-, climate- and life-history-based F_ST_ analyses (**Fig. S3**), have F_ST_^time^ values close to the mean or among the 10% lowest values of their respective control genes, suggesting that their differentiation following geography and climate might have been consistent overtime and already present historically (see also **Fig. S6d**). In contrast, the cell-cycle gene *CYCD5;1* (AT4G37630, affects divisions in the stomatal lineage; (Han et al. 2018)), *SCAP1* (AT5G65590, a transcription factor in the stomatal lineage; (Castorina et al. 2016)) and *STOMAGEN* are highly differentiated overtime, with F_ST_^time^ among the 10% highest values of their background genes (for changes in Tajima’s D between historical and modern paired samples, see **Fig. S6b,c**). It is crucial to note that the observed pattern may reflect dataset-inherent differences that are difficult to account for, such as the excess of low-frequency variants that is driving the modern data’s negative mean Tajima’s D (**Fig. 1e**). We aimed to address this by including only high coverage SNPs shared by both datasets addresses. However, this stringent filtering may also reduce the analyses’ sensitivity, especially as the final paired dataset is small (126 sample pairs). Differences in population history and sampling-location specific climate change that remain despite the geographic sample pairing add further noise to these analyses, as does the broad time window of samples classified as “historical” (1817-2002). Nevertheless, some genes seem to show indications of change across temporal and geographic gradients. Their potential connection to climate change adaptation will however require further analyses and experimental verification.

#### Text S2. Stomatal density score correlation with latitude

*A. thaliana’s* extensive population structure complicates disentangling demographic from adaptive genetic signatures (1001 Genomes Consortium 2016). We attempt to address this in our analyses of stomatal density score correlations with latitude by permuting genetic and phenotypic signals (see main text and **Fig. S4d,e**), and formally correcting for population structure using a genetic relationship matrix (see below).

We used the published 1001G SNP-matrix to decompose genetic relationships between the 1325 modern accessions with a principal component analysis (PCA; Plink v1.9, (Purcell et al. 2007; Chang et al. 2015)). Modeling *stomatal density score ~ latitude* confirms the explanatory power of samples’ latitude of origin for the stomatal density score (p-value_lat_ = 3.74 10^−13^, R^2^_lat_ = 3.88 10^−2^, **Table S11**). We also calculated latitude correlations for 100 random independent subsets of 126 modern samples. These subsets retain the correlation pattern (all slopes positive, and significant in 69 of 100 samplings; **Fig. S4a, b**, **Table S11**), indicating that the trend is unlikely due to geographic sampling bias.

When correcting for population structure, addition of the first three PCs marginally increases the percentage of variance explained by the model (*stomatal density score ~ latitude + PC1 + PC2 + PC3;* R^2^_lat_ = 0.039, p-value_lat_ = 3.741 10^−13^, R^2^_lat+PC123_ = 0.064, p-value_lat+PC123_ = 3.001 10^−16^, **Table S11**). However, latitude remains the single significant coefficient (p-value_lat_= 1.42 10^−15^, p-value_PC1_= 0.119, p-value_PC2_ = 0.792, p-value_PC3_ = 0.914). This is consistent also in the subset of 126 modern samples that are paired with a historical counterpart (p-value_lat_ = 0.012, p-value_PC1_ = 0.515, p-value_PC2_= 0.671, p-value_PC3_= 0.244), as well as in the majority of 100 random subsets of 124 modern samples (11/100 subsets with only p_lat_ < p_pc1_, 25/100 plat < p_pc1_ ***** p_c2_, 42/100 p < p_pc1_ p_c2_ p_c3_, i.e. a total of 78/100 with at least p_lat_ < p_pc1_ of these, 51/100 have only latitude as a significant coefficient with p < 0.05, PC1 is significant in 18/100, of which 14 also have latitude significant, with a lower p-value for latitude than any PC in all cases). The correlation between the stomatal density score and latitude thus seems to go beyond reflecting population structure alone.

#### Text S3. Extended analyses of stomatal density trends over time and with climate

The species’ demographic history and thus the geographic composition of the sample pairs may be contributing to the shifts in density score (delta_score_) we observe over time and associated with specific climate parameters (**Fig. 3, S7**; see also **Text S2**).

To assess the effect of geographic composition of sample pools on our analyses, we generated different subsets of the original 126 paired samples, excluding either all samples from the USA (retaining samples from above longitude -50°N; remaining n_pairs_= 104), from the USA and Spain (latitude > 44° N, n_pairs_ = 78), all samples from Scandinavia (latitude < 54°N, n_pairs_ = 98), or all sample pairs where the collection date of one (modern) sample is unclear (n_pairs_= 92). In all sets, stomatal density decreased in modern samples, but the trend becomes non-significant when excluding samples from latitudes lower than 44°N (USA, Spain; p-value_noUS_ = 0.368, p-value_noUS___noSpain_ = 0.113; **Table S7**, **S8**). While this could suggest that the source of the temporal signal is in the US and Spain, we refrain from jumping to this conclusion given that these large scale sample exclusions could simply cause a drop in statistical power, as non-US/non-Spain samples also display a trend of decreased stomatal score, which is however not significant for these samples alone (**Table S7**, **S8**).

To take a more continuous approach to studying stomata decrease over time, rather than subsetting the data in regions, we fit the first three genomic principal components (PC) in a linear regression model (*delta_score_ ~ mean + latitude + PC_1_ + PC_2_ + PC_3_*), where each predictor variable is mean-centered. In this model, the mean intercept was negative and significant (p-value = 1.51 10^−14^), indicating there is an average decrease in stomatal density score after accounting for confounders (**Table 11**).

We also tested whether the association we find between decreasing stomatal density and increased precipitation may be biased by population structure. Again, we find that the association is significant with p < 0.05 in all subsets except those excluding samples from the USA (*delta_score_ ~ precipitation_directionaiiy_;* see **Table S9**, and testing with Fisher’s Exact test in **Table S10** and **Fig. S7e,f**), and increased precipitation is the coefficient with the strongest effect on delta_score_ (estimate_precipitation_ = -1.074).

In addition, we test for the effect of population structure within all subsets by using PCs as described in **Text S2**. Inclusion of any of the first three PCs makes the model non-significant across all subsets (*deltas_C_or_e_ ~ precipitation_d_ir_e_c_t_ionality* p-value = 0.045; *deltascore ~ precipitation_d_ir_e_c_t_ionality + PCx* p-value > 0.05, see also **Table S9**), and even in subsets where the model and effect of increased precipitation are not significant with p-value < 0.05, increased precipitation consistently has a lower p-value than any of the PCs.

Thus, while both geographic sampling distribution and PCs as “formal” indicators of population structure do have an effect on the observed decrease in stomatal density over time, and on the association of this trend with increased precipitation, both trends remain consistent (if not always significant) across our validations.

##### Note on physiologically-relevant temperature values

Even combinations of precipitation and temperature change to some extent conformed to previous studies, where stomatal density did not change significantly from historical to modern samples when experiencing increased maximum temperature and decreased water availability together (odds_t_max_hi_g_h_pptj¤w = 0.588, ptmax_high_ppt_low = 0.279; **Fig. S7e,f**, (Vile et al. 2012)). For increased temperature alone however, unlike published experiments (e.g. (Vile et al. 2012; Lau et al. 2018; Crawford et al. 2012)), we only detected a weak tendency for stomatal density decrease. Such weak signals for temperature-related responses (see **Table S10**) may also result from the relatively large range of moderate temperature increases measured across our sample sites (0.7-3°C, mean 2.2°C; modeled per site using monthly maximum °C and calculating the °C increase over the recorded time frame (1958-2017) with the product of 59 years times the slope of each sites’ linear regression model, only considering sites with slopes of p-values < 0.01 after Benjamini Hochberg correction). In contrast, experimentally observed density decreases, which guided our intuition of stomatal change, were conducted in laboratory growth conditions by increasing temperatures by 6-10°C (22 to 28°C in (Crawford et al. 2012; Lau et al. 2018), 20 to 30°C in (Vile et al. 2012)). In

26 agreement with our observations, others have reported diverse effects of temperature (and CO_2_) on stomatal density (stomatal density increases, decreases and no effects), while water deficit elicited a consistent response also across species (see meta-analysis (Yan, Zhong, and Shangguan 2017)).

In addition, overall climate change and plant response trends over time in the present datasets may be weakened or obfuscated by the relatively short timeframe of available high-resolution historical global climate data (1958-2017, (Abatzoglou et al. 2018)), and the generalization of all paired samples from herbaria, collected between 1817 and 1957, as “historical” and all of the 1001G accessions, collected between 1992 and 2012, as “modern”. Nevertheless, we identify trends that correspond well both with experimental and historical observations of stomatal density responses to the climate variation connected with global change.

#### Text S4. Stomatal density versus stomatal index measurements

We chose to generate a polygenic score based on stomatal density rather than stomatal index (SI), i.e. the ratio of stomata to the total number of epidermal cells, as much fewer experimental studies have detailed the effects of loss of stomatal development genes on SI, while stomatal density is routinely measured. Notably, studying SI might to an extent alleviate some of the limitations of predicting stomatal density change accurately. While stomatal density reflects the leaves’ physiological capacity and varies greatly with cell or leaf size changes, SI is independent of the latter and a direct indicator of the cellular and developmental processes generating stomata and leaf growth. As environmental changes may affect both in different ways, complementary studies can expand our understanding of which developmental pathways of plasticity and growth change affect SI, and how this modulates stomatal density and thus gas exchange. Studying SI therefore represents a promising avenue for future studies to better understand the nature of changes in stomatal development following climate change.

### Supplemental Tables

Supplementary tables available at subsequent link: https://docs.google.com/spreadsheets/d/1TaKYvIKgS5DpuW_iuv9NRXMlsuuyciFtiMNIP-4-n6A/edit?usp=sharing

**Table S1.** *Historical sample information*

**Table S2.** *Stomatal development genes.*

**Table S3.** *Presence of different SNP categories in stomatal core genes.*

**Table S4.** *Stomatal genes used for PGS.*

**Table S5.** *Correlations of whole-genome and stomata gene PGS versus measured stomatal density and stomatal density score.*

**Table S6.** *Historical and modern sample pairs.*

**Table S7.** *Stomatal density changes across sample subsets (Student’s T-test).*

**Table S8.** *Stomatal density correlation with latitude and change across sample subsets.*

**Table S9.** *Stomatal density change correlation with climate (Linear models).*

**Table S10.** *Stomatal density change correlation with climate (Fisher’s Exact test).*

**Table S11.** *Stomatal density score correlation with latitude with population structure correction.*

**Fig. S1:**
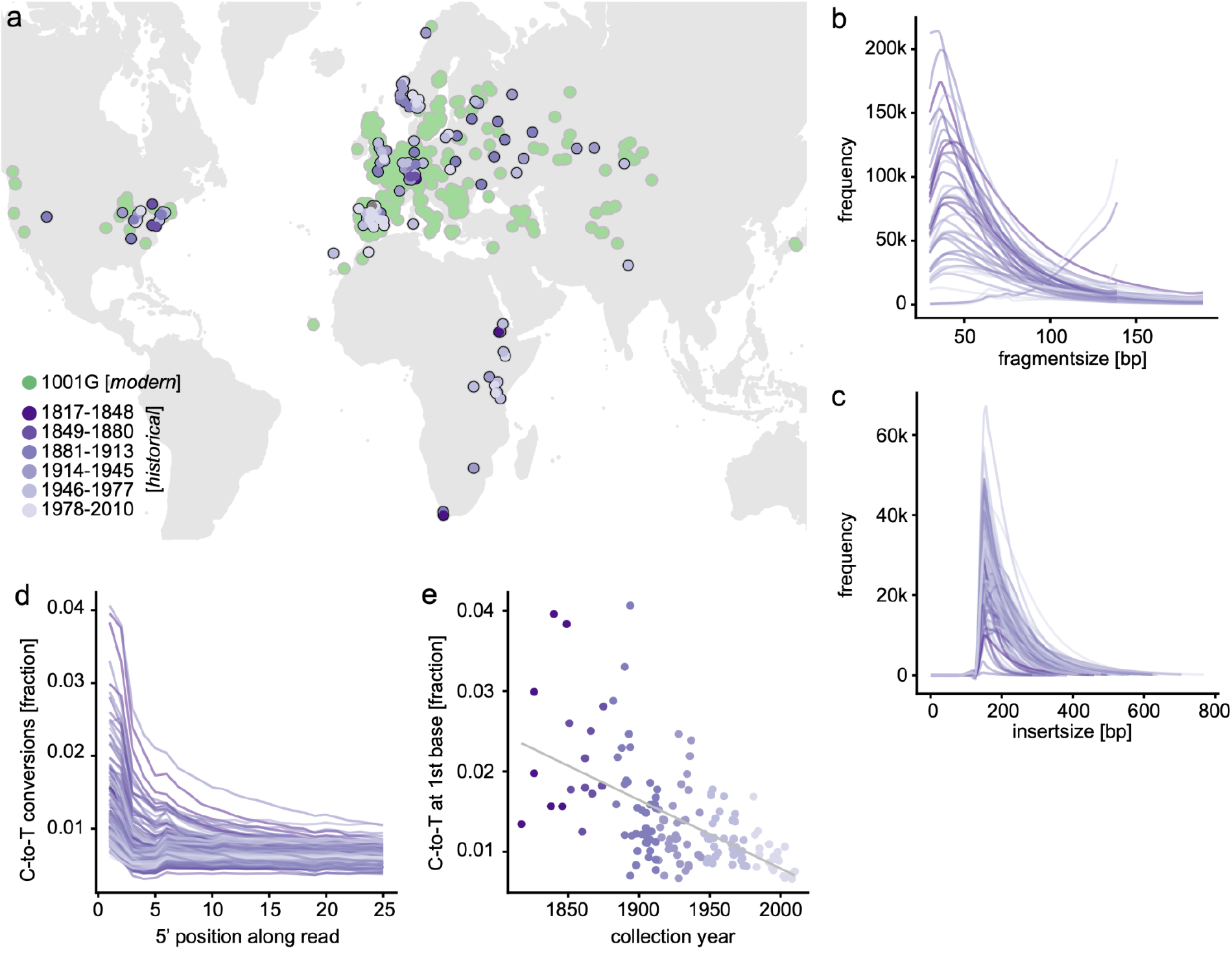
Worldwide samples and historical sample authentication. (**a**) Map of geographic origin of sequenced historical specimens, colored by sample age (Lopez et al. 2022). Darker purple shades indicate older samples, green visualizes geographic origin of the published 1001 A. thaliana genomes (1001 Genomes Consortium 2016). (**b**, **c**) Sequenced DNA fragments of historical samples, (**b**) merged sequencing reads for unsheared historical DNA and (**c**) insert sizes of sheared and unmerged DNA fragments. (**d**) Fraction of C-to-T converted base-pairs along sequencing reads, and the (**e**) correlation of the fraction of C-to-T conversion at the first base of a read with the age of the respective historical herbarium specimen (one-sided Pearson’s correlation test, correlation coefficient r = −0.589, p = 1.693 10^−15^).

**Fig. S2:**
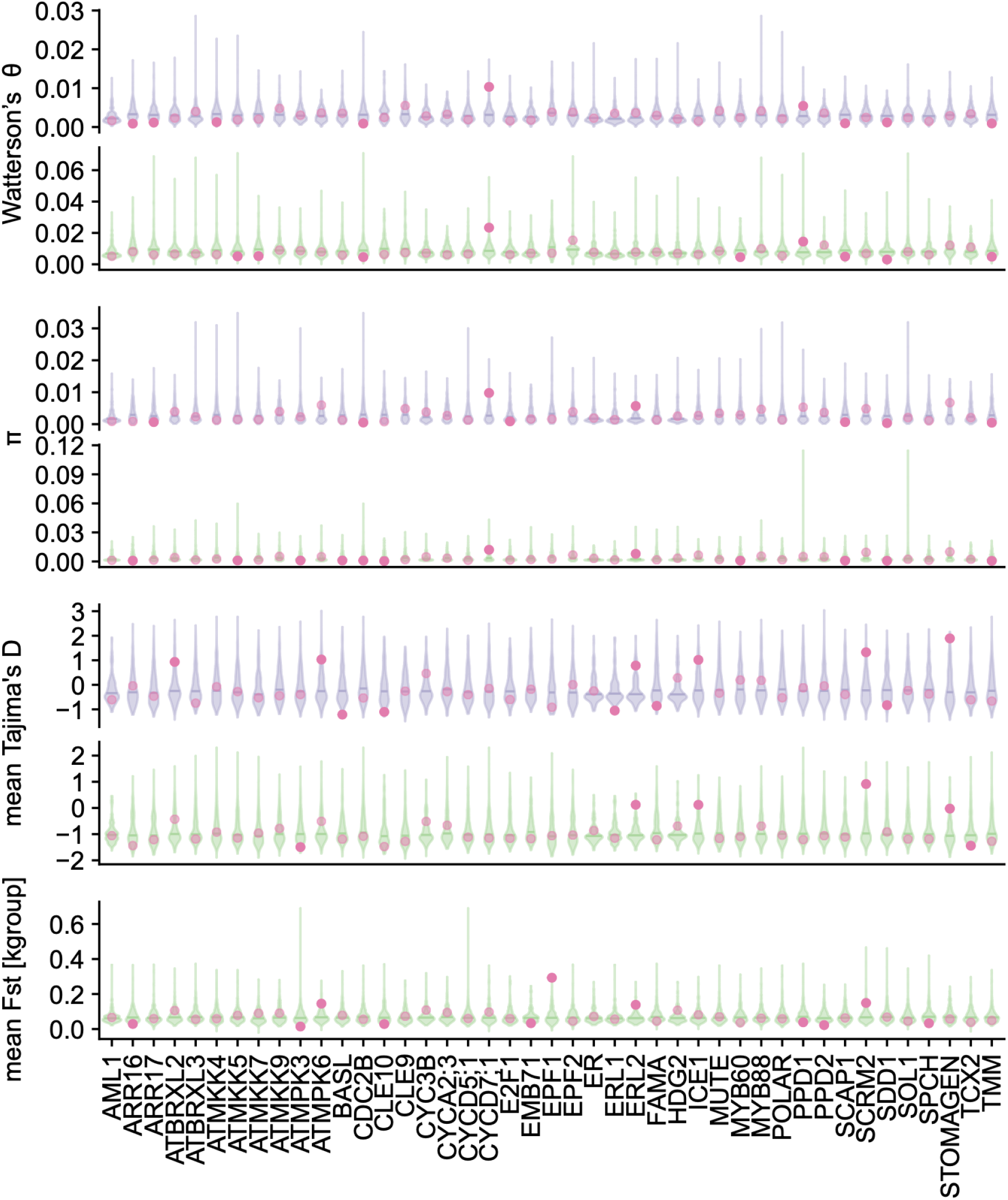
Per-gene values of diversity statistics. From top to bottom, per-gene mean values for Watterson’s θ, nucleotide diversity π and Tajima’s D (assigned to SNPs from 100-bp-window calculations) for historical and modern samples, and for modern samples only F_ST_ (for the k-groups from (Exposito-Alonso, Vasseur, et al. 2018)). Distribution of control genes of comparable length in violin plots with horizontal line marking the 0.5 percentile, purple for historical, green for modern dataset. Transparent magenta points indicate the gene mean, solid magenta points indicate that the gene mean is among the 10% lowest or highest values of the control distribution.

**Fig. S3:**
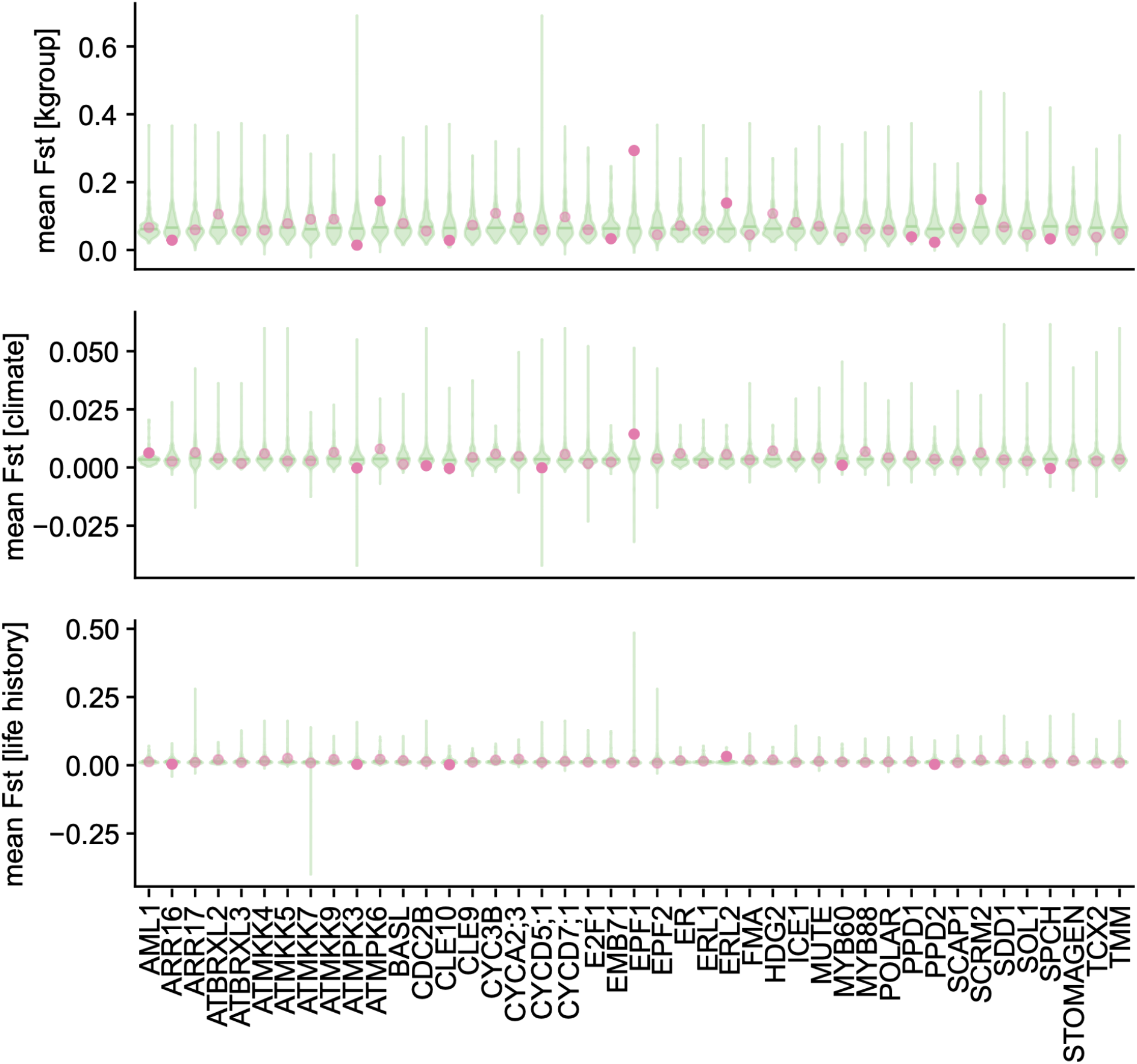
Fixation index for different population delineations. Mean per-gene F_S_t across the modern samples (green), calculated for groups defined by (from top to bottom) whole genome genetic variation (‘kgroups? as defined in (Exposito-Alonso, Vasseur, et al. 2018); plot identical to Fig. S2, reproduced here to facilitate comparisons), climate variation (precipitation and temperature seasonality as defined in BIO4 and BIO15, (Hijmans, Cameron, and Parra 2005)) and life history traits (as defined in (Exposito-Alonso 2020)). Distribution of control genes of comparable length in violin plots with horizontal line marking the 0.5 percentile. Transparent magenta points indicate the gene mean, solid magenta points indicate that the gene mean is among the 10% lowest or highest values of the control distribution.

**Fig. S4:**
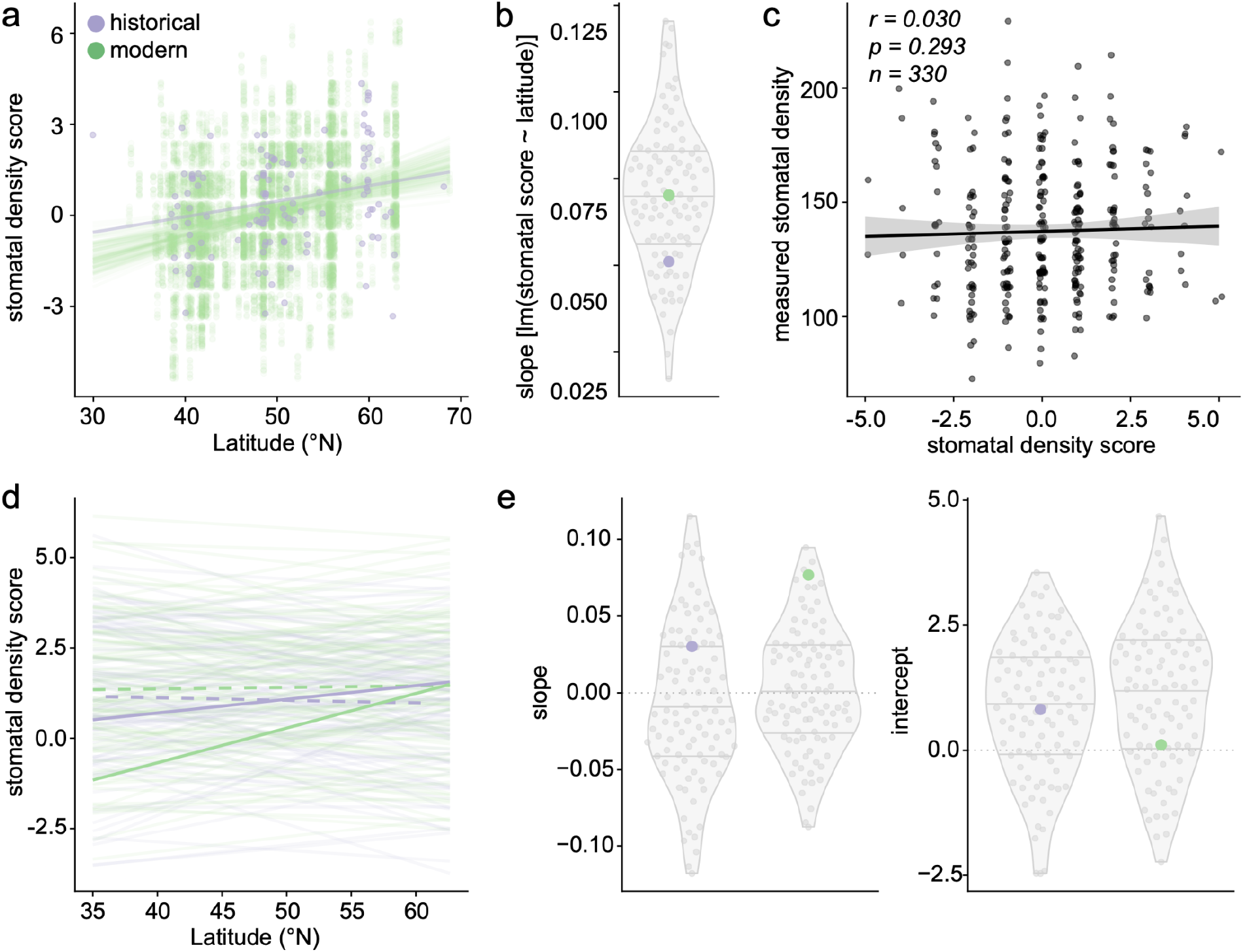
Latitude patterns validate biological relevance of stomatal density score. (**a**) Stomatal density score correlations with latitude at samples’ geographic origins. Historical samples (124, purple) and 100 resamplings (without replacement) of 124 modern samples (green), drawn from the 1001G data, with linear regressions indicating increasing stomatal density with increasing latitude, as also observed for experimentally measured stomatal densities (Dittberner et al. 2018). (**b**) Violin plot of slopes from regressions in (**a**), with 69/100 significant with p < 0.05, and 100/100 positive. (**c**) Comparison of the density score with measured stomatal densities (positive but not significant trend, linear regression slope = 0.447, one-sided Pearson’s correlation test, p-value = 0.293, correlation coefficient r = 0.030, data from (Dittberner et al. 2018)). (**d**) Linear regressions of the same parameters as in (**a**), over 100 permutations of genes and their effects on stomatal density (increase/decrease), for historical (purple) and modern data (green). Solid lines for original data (statistics see **Fig. 2d**, **Table S11**), transparent lines for permutations, dashed lines for mean of permutations. Permutation means are not significantly correlated with latitude (one-sided Perason’s correlation test, p_mod___mean_ = 0.445, p_hist___mean_ = 0.134, correlation coefficient r_mod___mean_ = 0.067, r_hist___mean_ = −0.134). Linear regressions of the permuted density score with latitude are mostly not significant, and randomly positive or negative (9/100 modern and 20/100 historical significant slopes with p < 0.05; positive slopes in 45/100 historical and 51/100 modern regressions). (**e**) Slopes (left) and intercepts (right) of the permutations shown in (**d**), with permutations in gray, and original data in purple (historical) and green (modern). Intercepts were calculated from linear regression models, using the samples’ median latitude to avoid extrapolation beyond the geographic space covered by the samples. Density differences over time in the density score-latitude association indicate a potential density decrease in modern samples (density score ~ latitude; y-intercept_modern_ = 0.121, y-intercept_historical_ = 0.799), which is confirmed by a linear regression model integrating latitude and dataset identity (historical vs modern; density score ~ latitude + dataset; p-value_lat_ = 0, p-value_dataset_ = 0, p-value_model_ = 7.732 10 ^6^). Dotted horizontal lines mark slope /intercept of 0, corresponding to the expectation given full latitude independence of the permuted phenotypes. The permuted samples’ positive average intercept indicates a slight positive bias of the stomatal density score, possibly due to the higher number of genes positively (14) vs negatively (10) affecting stomatal density used for the score’s calculation. Analyses exclude samples from North America and the African continent.

**Fig. S5:**
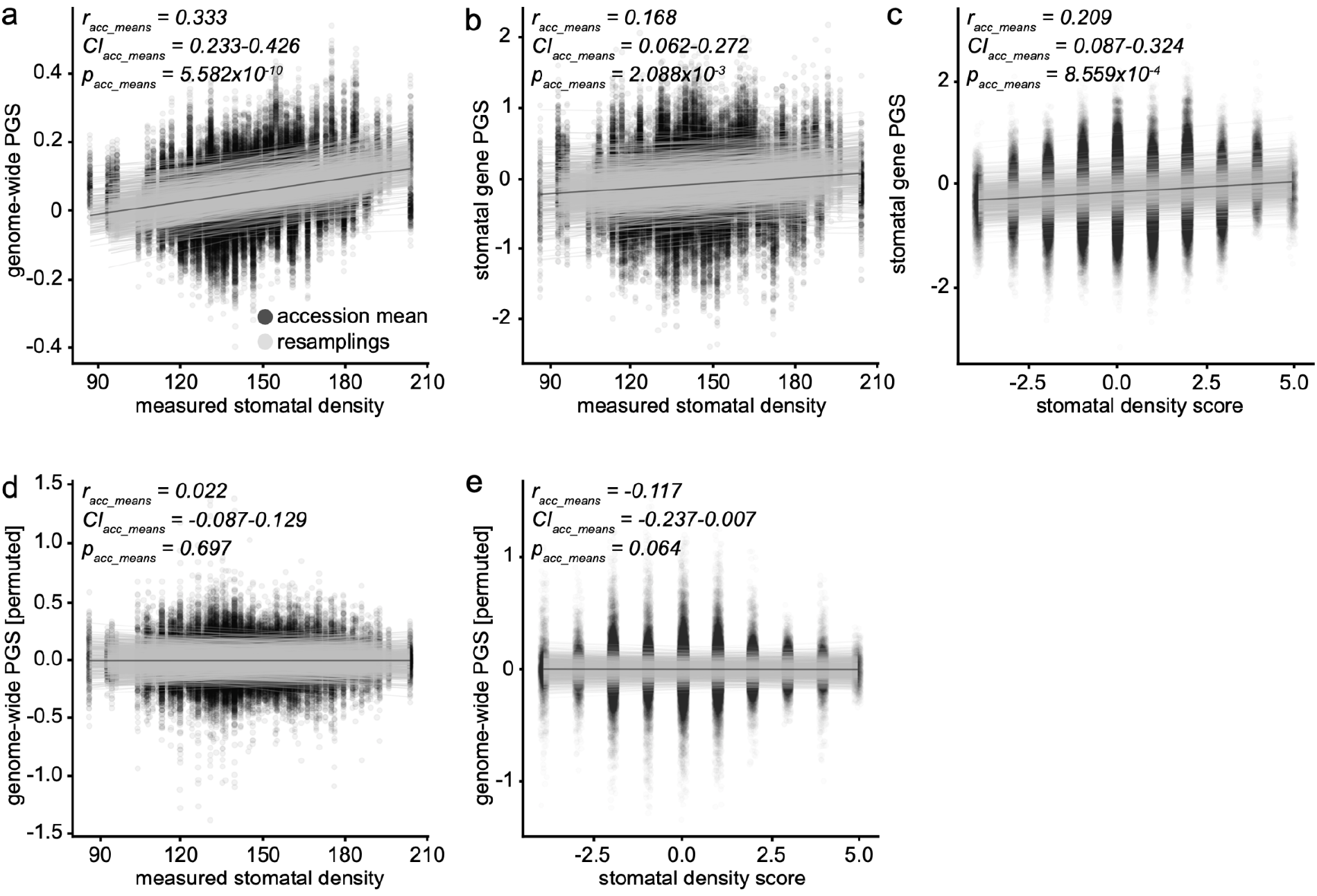
PGS correlates with measured stomatal density and stomatal density score. (**a**) Genome-wide PGS correlates with experimentally measured stomatal densities (Dittberner et al. 2018). One-sided Pearson’s correlation test, positive correlation in 1000/1000 re-trainings, 745/1000 significant one-sided Pearson’s correlation tests with p<0.05, correlation coefficient r_min-max_ = 0.010−0.561; r_median_ = 0.304. (**b**) PGS generated only based on 43 stomatal development genes correlates with experimentally measured stomatal densities. One-sided Pearson’s correlation test, positive correlation in 888/1000 re-trainings, 150/1000 significant one-sided Pearson’s correlation tests with p<0.05, correlation coefficient r_min___max_ = −0.209−0.418, r_median_ = 0.133. (**c**) Correlation trend between stomatal gene-based PGS and the stomatal density score. One-sided Pearson’s correlation test, positive correlation in 989/1000 re-trainings, 737/1000 significant one-sided Pearson’s correlation tests with p<0.05, correlation coefficient r_median_ = 0.16. (**d**) Genome-wide PGS based on a training-set with permuted genotype-phenotype associations. One-sided Pearson’s correlation test, positive correlation in 485/1000 re-trainings, 112/1000 significant one-sided Pearson’s correlation tests with p<0.05, correlation coefficient r_median_ = −0.008. (**e**) Stomatal gene-based permuted PGS. One-sided Pearson’s correlation test, positive correlation in 479/1000 re-trainings, 367/1000 significant one-sided Pearson’s correlation tests with p<0.05, correlation coefficient r_median_= −0.007. See also **Table S5**.

**Fig. S6:**
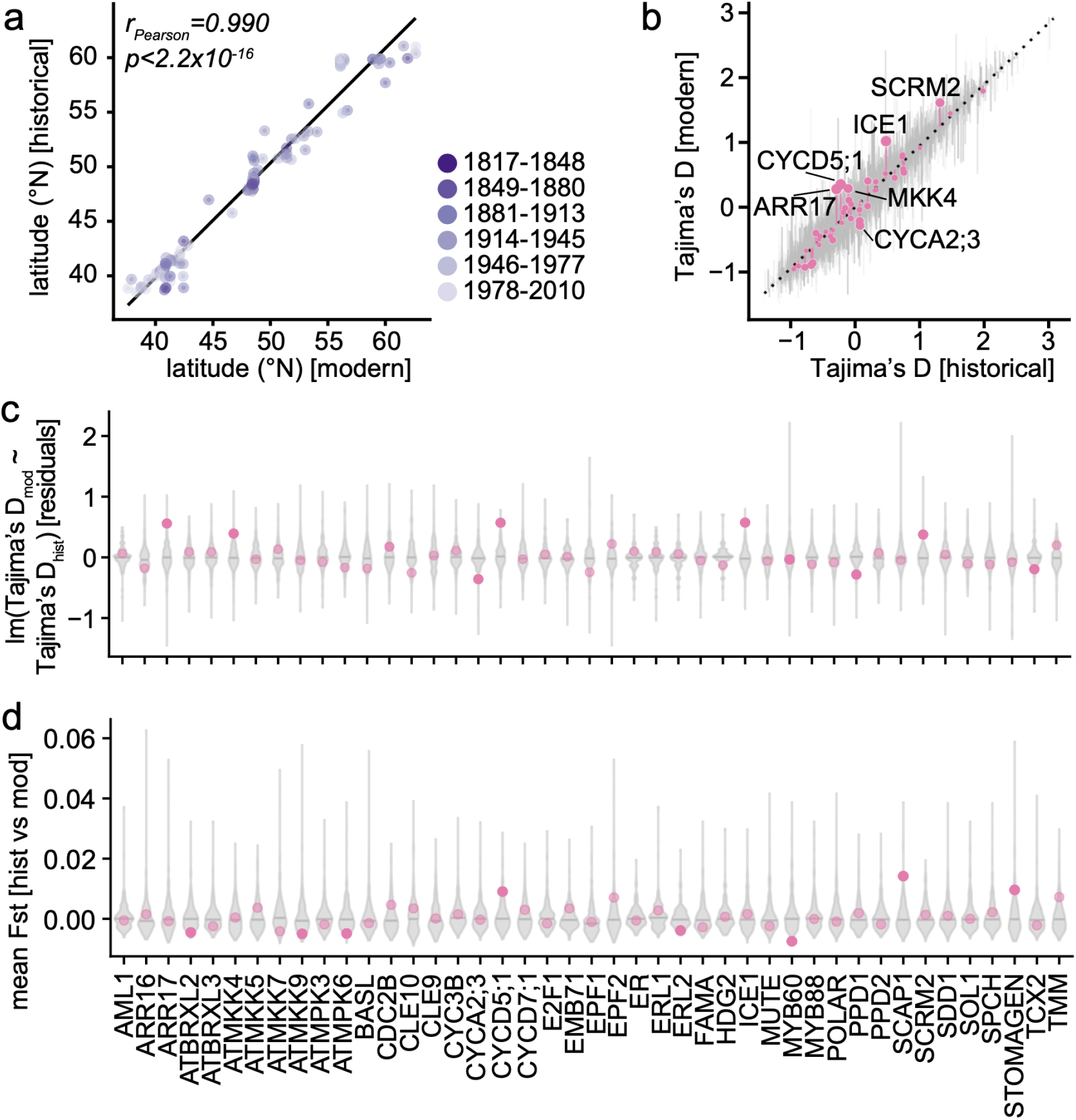
Temporal change across historical and modern sample pairs. (**a**) Geographical origin-based sample pairing reduces biases in sample distribution between historical and modern datasets, as indicated by the regression of paired samples’ latitudes of origin (one-sided Pearson’s correlation test, correlation coefficient r = 0.990, p < 2.2 10^−16^). (**b**) Comparison of Tajima’s D values between paired historical and modern datasets, displayed as residuals of a linear regression Tajima’s D_mod_ ~ Tajima’s D_hist_ (dotted line). Background control genes’ Tajima’s D in gray, and stomatal genes’ Tajima’s D in magenta. Circle-size for each stomatal gene indicating the magnitude of the gene residuals’ deviation from the regression. (**c**) Residuals of the linear regression model as in (**b**), per stomata gene and its specific control set. Distribution of control genes of comparable length in violin plots with horizontal line marking the 0.5 percentile. Transparent magenta points indicate the gene mean, solid magenta points indicate that the gene mean is among the 10% lowest or highest values of the control distribution. (**d**) Change in stomatal genes over time, as measured by F_ST_^tme^ between paired historical and modern sample groups.

**Fig. S7:**
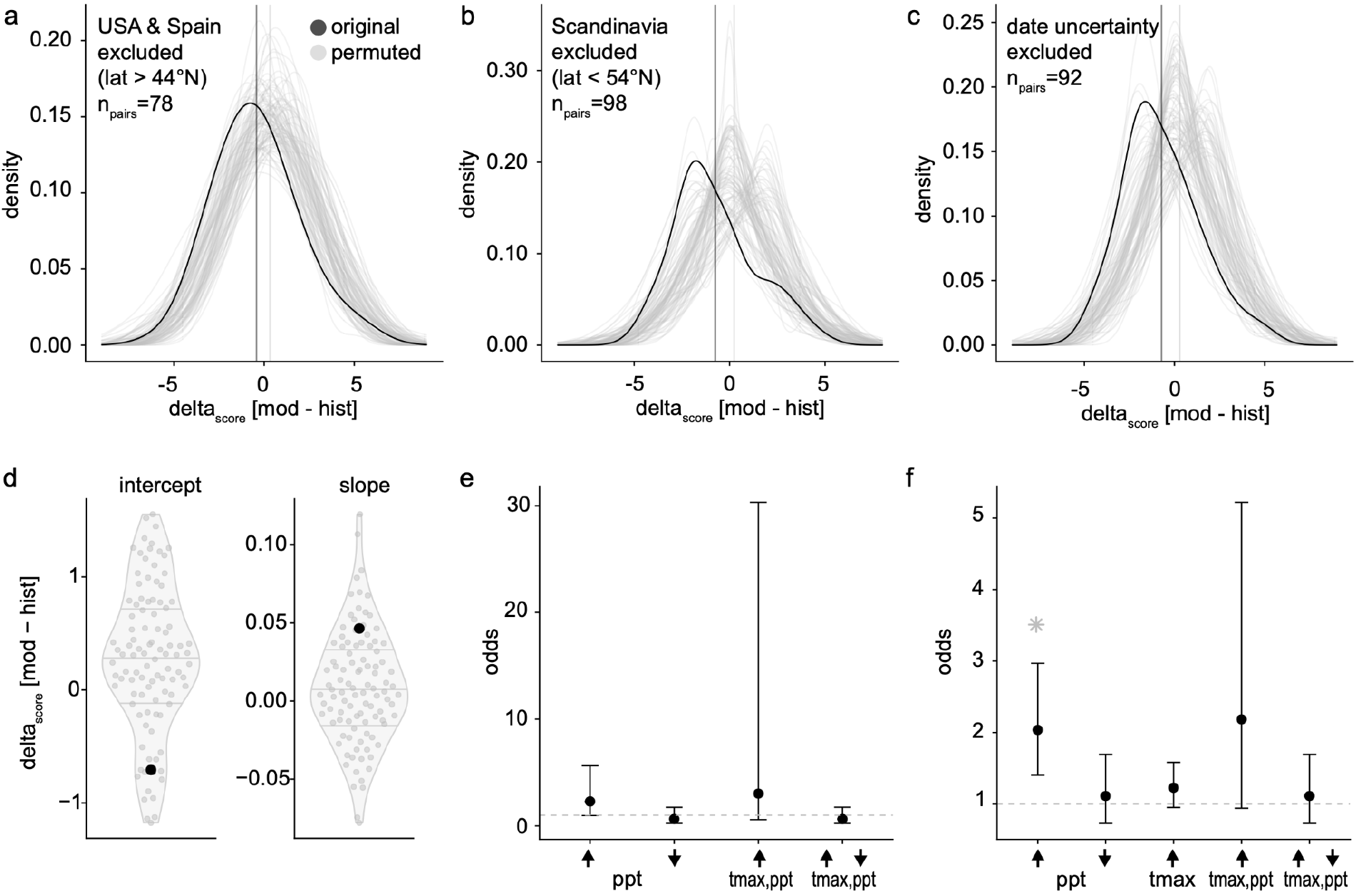
Stomatal density score decreases over time. (**a-c**) Distributions of per sample-pair calculated difference in stomatal density scores (delta_score_) for original data (black) and 100 permutations (gray) of genes and their assigned effect (decrease/increase) on stomatal density. Density distribution means are marked by solid black and gray vertical lines. See also **Table S7**, **S8**. (**a**) Dataset including only samples from latitudes >44°N, i.e. excluding the USA and Spain; mean delta_score_^noUS^-^noSpain^ = −0.745, smaller than 90 of 100 permutations, Wilcoxon signed rank test, p < 2.2 10^−16^. (**b**) Including only samples from latitudes <54°N, i.e. excluding Scandinavia; mean delta_score_^noScandinavia^ = −0.423, smaller than 93 of 100 permutations, Wilcoxon signed rank test, p < 2.2 10-^16^. (**c**) Excluding samples with uncertain collection year; mean delta_score_^noUncertainty^ = −0.717, smaller than 93 of 100 permutations, Wilcoxon signed rank test, p < 2.2 10^−16^. (**d**) Difference in slope and intercept between individual regressions of the stomatal density score with latitude for the full set of historical and modern samples (black) and 100 phenotype permutations (gray), to reflect change over time as delta_score_ while accounting for remaining geographic biases. Slope and intercept were extracted from each regression and subtracted to calculate delta_score_[mod-hist]_slope_ and delta_score_[mod-hist]_intercept_, the latter based on the median latitude of each sample group to avoid extrapolation beyond the geographic range covered by samples. The intercept, indicative of the decrease in stomatal density score over time, in the original data is lower than in 89/100 permutations, hinting at a density score decrease. The change in slope is higher than in 86/100 permutations. (**e**) Odds of stomatal density decrease over time, calculated on subsets of historical and modern sample pairs whose geographic origin has experienced a significant increase or decrease in precipitation (left), a significant increase in maximum temperature and precipitation (center), or a significant increase in maximum temperature combined with a decrease in precipitation (right) between 1958 and 2017. (**f**) Fisher’s Exact Test odds of modern samples containing more SNPs that decrease stomatal density than historical samples under specific environmental conditions, calculated independent of historical and modern sample pairings. From left to right, for all geographic locations (not necessarily paired) with increased precipitation, decreased precipitation, higher temperature, or higher temperature and precipitation from 1958 to 2017 (Fisher’s Exact test; asterisk indicates p < 0.001; see **Table S10**). Analyses include samples from North America and exclude samples from the African continent.

